# Genome-wide variability in recombination activity is associated with meiotic chromatin organization

**DOI:** 10.1101/2021.01.06.425599

**Authors:** Xiaofan Jin, Geoff Fudenberg, Katherine S. Pollard

**Affiliations:** Gladstone Institutes, San Francisco, USA; University of California San Francisco, San Francisco, USA; Chan-Zuckerberg Biohub, San Francisco, USA

## Abstract

**Background:** Recombination enables reciprocal exchange of genomic information between parental chromosomes and successful segregation of homologous chromosomes during meiosis. Errors in this process lead to negative health outcomes, while variability in recombination rate affects genome evolution. In mammals, most crossovers occur in hotspots defined by PRDM9 motifs, though PRDM9 binding sites are not all equally hot. We hypothesize that dynamic patterns of meiotic genome folding are linked to recombination activity.

**Results:** We apply an integrative bioinformatics approach to analyze how three-dimensional (3D) chromosomal organization during meiosis relates to rates of double-strand-break (DSB) and crossover formation at PRDM9 hotspots. We show that active, spatially accessible genomic regions during meiotic prophase are associated with DSB-favoured hotspots, which further adopt a transient locally active configuration in early prophase. Conversely, crossover formation is depleted among DSBs in spatially accessible regions during meiotic prophase, particularly within gene bodies. We also find evidence that active chromatin regions have smaller average loop sizes in mammalian meiosis. Collectively, these findings establish that differences in chromatin architecture along chromosomal axes are associated with variable recombination activity.

**Conclusions:** We propose an updated framework describing how 3D organization of brush-loop chromosomes during meiosis may modulate recombination.

## Background

The formation of crossovers during meiotic recombination is a highly orchestrated process, enhancing genetic diversity by allowing reciprocal exchange of genomic information to occur between parental chromosomes. Crossover formation also ensures proper segregation of homologous chromosomes[1], and errors in this process lead to chromosomal abnormalities such as aneuploidy which are associated with negative health outcomes[2, 3].

In mammals, crossovers are highly enriched (100 fold) in discrete ~1-2 kilobase (kb) sites along the genome, termed recombination hotspots[4]. Hotspot locations are in large part determined by binding of PRDM9, a meiosis-specific zinc-finger protein that marks genomic loci as candidates for recombination[5, 6, 7]. PRDM9 is thought to have at least two distinct modes of binding[8]. Class 1 binding results from motif-recognition of the PRDM9 zinc finger array, and marks genomic loci as candidates for recombination (i.e., hotspots). Class 2 PRDM9 binding occurs away from hotspots, and is proposed to represent tethering of hotspots to DSB machinery.

While hotspot initiation is dependent on PRDM9, subsequent DSB and crossover formation are highly stochastic: a mammalian chromosome may harbour hundreds of PRDM9 binding sites, but during a typical meiotic cycle only 10-20 double stranded breaks (DSBs) are catalyzed[9]. Out of these DSBs, most are repaired as non-crossover conversion events and only 1-2 per chromosome are chosen as sites for crossover[10, 11]. Local chromatin features such as GC content, histone modification and cofactor binding are known to impact DSB formation at hotspots[12], while nucleosome occupancy, GC content and chromosomal position are associated with crossover formation[13]. Still, a full understanding of why certain hotspots are favoured to form DSBs and crossovers remains undetermined.

Here we investigate how 3D genome folding relates to PRDM9 binding, DSBs, and crossover formation in mice. In particular, our work makes use of recent Hi-C datasets measuring the 3D conformation of meiotic chromosomes[14, 15, 16, 17, 18], and extends their findings on recombination[14]. Meiotic chromosomes adopt a brush-loop conformation characterized by chromatin loops attached to a central axis[19]. While recombination hotspots are found within loops, DSB machinery – such as DNA-repair proteins – resides on the axis[20, 21, 22, 23]. This “tethered-loop/axis complex” model of recombination suggests that 3D chromosomal structure could place constraints on the recombination process. We apply computational analyses that integrate observations from multiple recent interphase and meiosis datasets measuring recombination activity and chromatin organization, including Hi-C. We find that distinct chromatin environments are favourable for DSB and crossover formation respectively, and develop a statistical model of associations between chromosomal structure and recombination activity. These analyses lead to an updated proposal regarding how meiotic events related to recombination are related to brush-loop chromosomal architecture.

## Results

To investigate the relationship between meiotic chromatin structure and recombination in the mouse genome, we integrate multiple published datasets (Fig. 1A) by uniformly mapping each dataset to a consistent set of 5kb bins across the autosomal chromosomes (492,557 bins genome-wide). Meiotic patterns are compared to counterparts from embryonic stem (ES) cells serving as an example of interphase chromatin organization[24, 25, 26], allowing the identification of meiosis-specific patterns.

**Figure 1:**
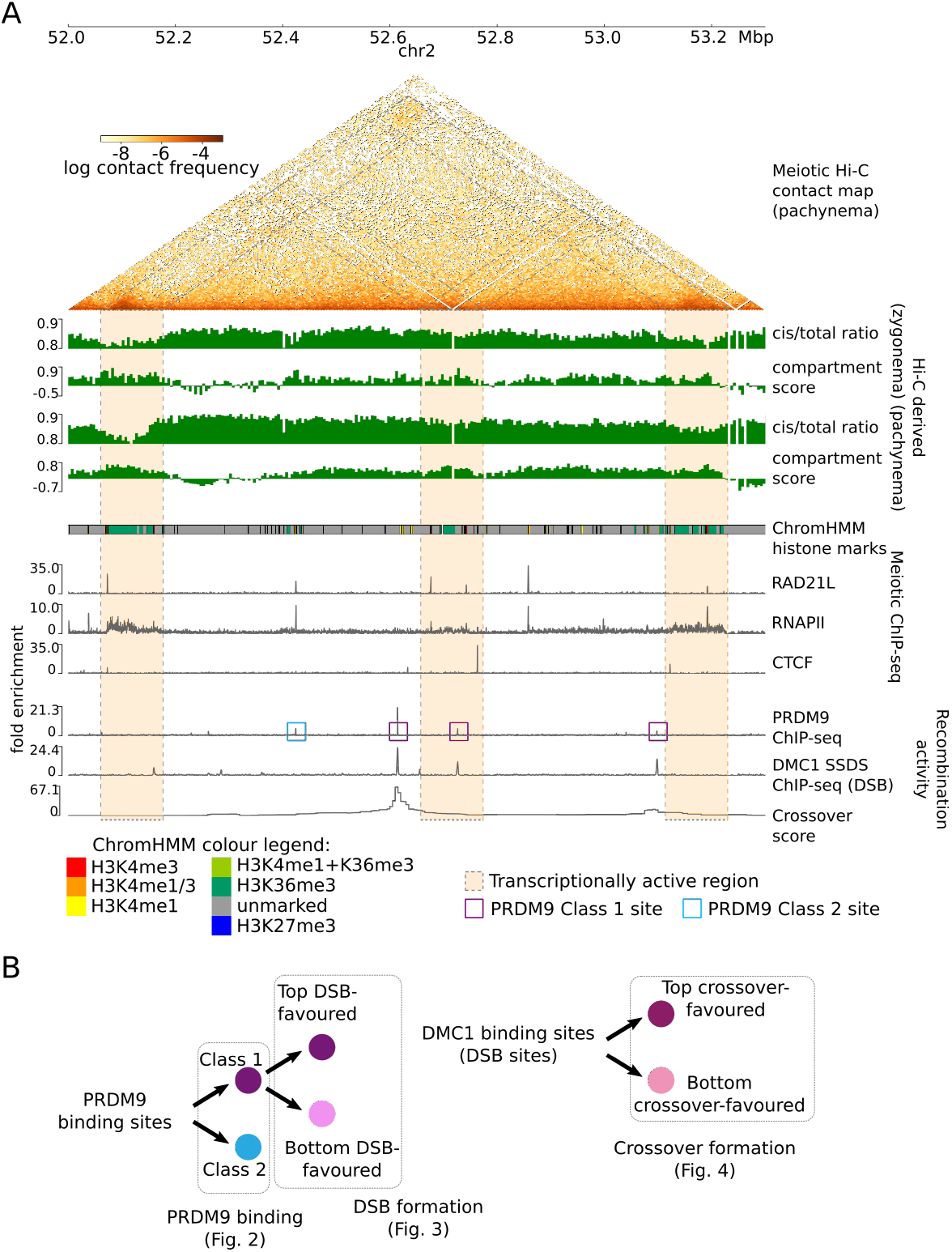
Multiple chromatin organization datasets are integrated with measurements of recombination activity at the levels of PRDM9 binding, DSB activity and crossover formation. **A:** Key datasets used in this study, shown in a browser view of a representative 1.3 megabase (Mb) region on mouse chromosome 2. Pachynema Hi-C contact frequencies are shown as a heatmap, in addition to Hi-C derived cis/total ratio and compartment score for zygonema and pachynema. Hi-C contact information is accompanied by epigenetic chromatin state information using chromHMM annotations of histone marks in mouse testis (see bottom left for colour legend), as well as meiotic ChIP-seq tracks of cohesin subunit RAD21L, RNAPII, and CTCF. Recombination activity measurements include ChIP-seq binding tracks of PRDM9, DMC1 (marking double stranded breaks), as well as crossover likelihood score derived from single sperm whole-genome sequencing. Several relationships to note in this region: (i) enriched Hi-C contacts between transcriptionally active regions during meiosis, highlighted in orange shaded boxes, (ii) colocalized meiotic cohesin occupancy, RNAPII and Class 2 PRDM9 binding, (iii) colocalized PRDM9 Class 1 binding, DSB formation and crossover likelihood, and (iv) locally depressed cis/total ratio and elevated compartment score at DSB sites in zygonema. Hi-C bins with missing data are ignored for visualization. **B:** Overview of recombination-related comparisons explored in this manuscript. PRDM9 binding sites are classified as Class 1 and 2 as previously described[8], and chromatin environments between these two classes are compared in Fig. 2. Subsequently, PRDM9 Class 1 sites favoured to form DSBs are compared against Class 1 sites disfavoured to form DSBs (Fig. 3). Finally, DSB sites favoured to form crossovers are compared against DSB sites disfavoured to form DSBs (Fig. 4)

We analyze meiotic chromosomal structure using Hi-C contact maps of mouse spermatocytes in the zygotene and pachytene stages of prophase I [14, 16, 17, 15], from which we derive genome-wide measures of meiotic chromatin 3D structure: cis/total ratio, A-B compartment scores, insulation scores and FIRE scores. We focus primarily on the first two measures, which display particularly interesting patterns related to recombination. Cis/total ratios quantify the fraction of contacts within versus between chromosomes. Low cis/total ratios are associated with increased spatial accessibility[27] (not to be confused with DNA accessibility associated with nucleosome occupancy). At a chromosome-wide level, lower cis/total ratios indicate a greater degree of chromosome territoriality[28]. A-B compartment scores quantify preferential interactions after removing the impact of genomic distance; high A-B compartment scores are typically associated with active, gene-rich chromatin (A-compartment) [29]. Meanwhile, insulation scores typically exhibit minima at domain boundaries[30], while high FIRE scores indicate regions with enriched interactions[31]. We supplemented these Hi-C metrics with 1D measurements of chromatin state, including chromHMM profiles from mouse testis[32] and occupancy patterns of CTCF, meiotic-specific cohesin subunit RAD21L [17] and RNA polymerase II (RNAPII)[33].

Alongside measurements of chromatin organization, we analyze PRDM9 binding measured by ChIP-seq[8], DSB activity based on DMC1 single-stranded DNA sequencing (SSDS) ChIP-seq[34], and crossover likelihood quantified in single-sperm genome sequencing datasets[35]. We explore differences in the 3D and local chromatin environment associated with **(i)** different classes of PRDM9 binding, **(ii)** variable likelihood in forming DSBs at PRDM9 binding sites, and **(iii)** variable rates of crossover selection at DSB sites (Fig. 1B), and integrate these findings with statistical models.

### Transient and locally active, spatially-accessible chromatin at PRDM9 Class 1 sites

We begin by exploring chromatin organization at PRDM9 binding sites, which mark candidate loci during leptonema for DSB formation – a prerequisite for crossover formation. We stratify our analyses by PRDM9 site type: PRDM9 Class 1 binding sites that occur at recombination hotspots, and Class 2 binding sites that occur away from recombination hotspots[8]. We focus on sites from the CAST *Prdm9* allele[8] – similar results were found with B6 allele, and we address differences where applicable (Supplements S1).

We confirm that PRDM9 Class 1 binding sites display increased DSBs and crossovers (Fig. 2A, 6421 genomic bins), and that they are found with reasonable frequency in both A and B-compartment bins (59%A– 41%B, Supplemental Fig. S1J). During zygonema, Class 1 sites are characterized by locally decreased cis/total ratio and elevated A/B-compartment scores (Fig. 2B) within a +/−50kb window. These transient zygonema-specific signals reflect active and spatially accessible chromatin respectively, and coincide with H3K4me3 trimethylation activity of PRDM9 at hotspots[6, 7, 5], which peaks in zygonema and fades in pachynema[36]. Averaged together, Class 1 sites exhibit a distinctive local Hi-C contact map specifically during zygonema (Fig. 2C). Note these meiotic Hi-C patterns are much weaker than signals in interphase such as TAD boundaries (Supplemental Fig. S2F), consistent with the general attenuation of TAD and compartment patterns in meiotic Hi-C datasets[14].

**Figure 2:**
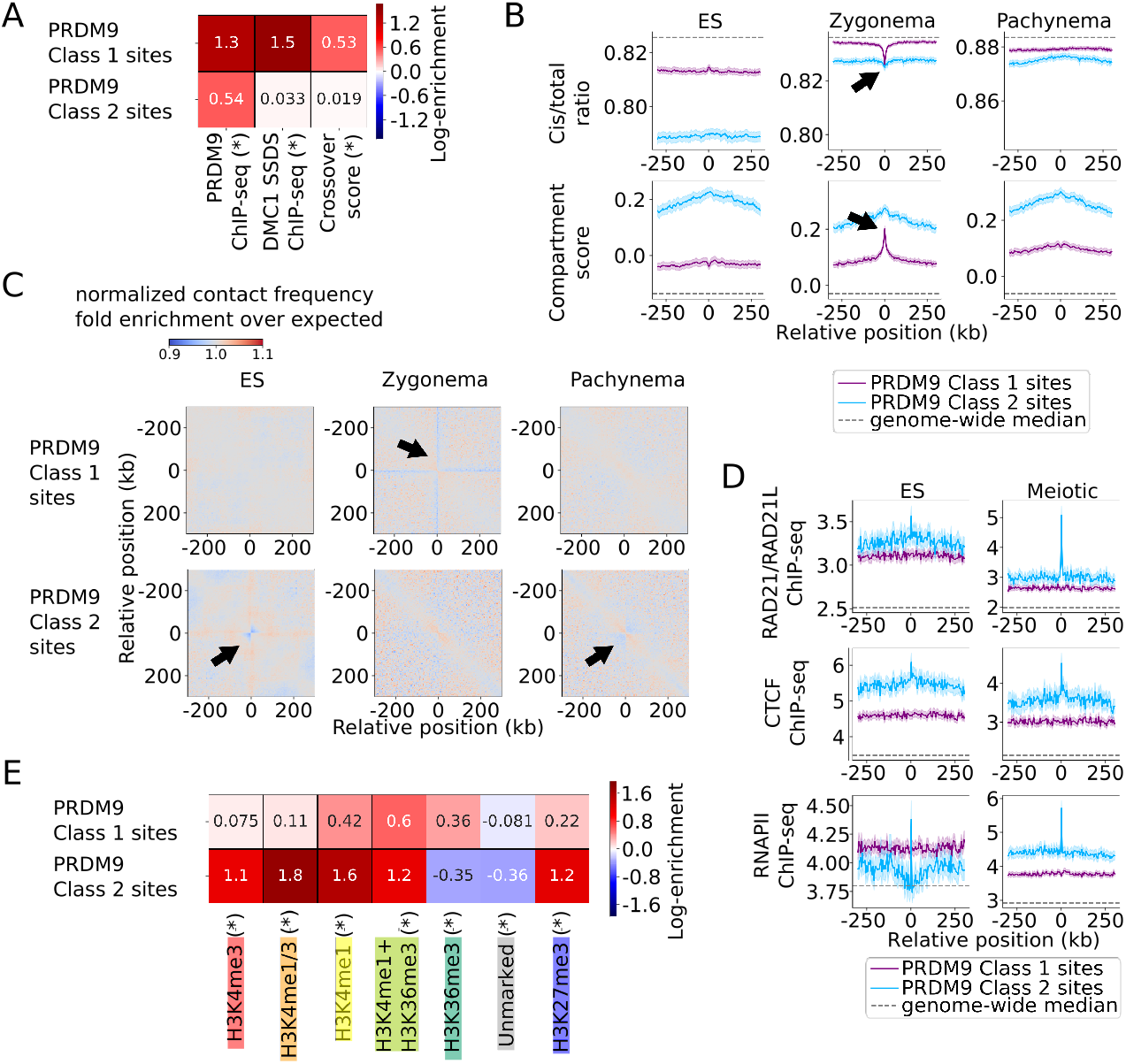
Distinct chromatin environments characterize PRDM9 Class 1 and Class 2 binding loci. **A:** PRDM9 ChIP-seq sites are classified into either Class 1 or Class 2 sites[22]. Class 1 sites represent recombination hotspots associated with increased DSB activity and crossover likelihood, while Class 2 sites are associated with later binding to recombination machinery and do not mark hotspots themselves. Heatmap shows log fold enrichment over 500 kb surrounding region, while (*) indicates Bonferroni-adjusted p<0.01 difference between PRDM9 Class 1 and Class 2 sites. **B:** Hi-C cis/total ratio (top) and compartment score (bottom), averaged across all PRDM9 Class 1 and 2 binding positions, calculated for ES, zygonema and pachynema datasets (columns). Shading represents 95% confidence intervals. Class 2 sites are associated with higher compartment score and lower cis/total ratio in general. Black arrows indicate zygomena-specific shifts toward active spatially accessible chromatin at PRDM9 binding sites. **C:** Normalized chromatin contact matrices, averaged across all Class 1 (top) and Class 2 (bottom) binding positions, for embryonic stem cell (ES), zygonema and pachynema Hi-C datasets (columns). Black arrows mark zygonema specific patterns in Class 1 sites, and weak boundary-like contact patterns at Class 2 sites. **D:** Cohesin subunit (top), CTCF (middle), and RNAPII (bottom) ChIP-seq tracks, averaged across all PRDM9 Class 1 and 2 binding positions, calculated for ES and meiotic datasets. Shading represents 95% confidence intervals. Class 2 sites exhibit overlap with cohesin, CTCF and RNAPII sites, particularly during meiosis. **E:** Overlap of chromHMM epigenetic state annotations with PRDM9 Class 1 and 2 binding sites indicates a strong enrichment of promoter and enhancer-like (H3K4me1/3, H3K27me3) histone marks in Class 2 sites, while Class 1 sites appear more evenly distributed across epigenetic states. Heatmap shows log fold enrichment over genomewide mean, while (*) indicates Bonferroni-adjusted p<0.01 difference between PRDM9 Class 1 and Class 2 sites.

### PRDM9 Class 2 binding specifically enriched in active chromatin

In contrast with Class 1 sites, PRDM9 Class 2 sites are not enriched for DSB or crossover activity and have a weaker PRDM9 ChIP-seq signal (Fig. 2A, across 2332 bins). Class 2 sites exhibit globally reduced cis/total ratio and elevated compartment scores (Fig. 2B). Unlike Class 1 sites, they are heavily biased to active A-compartment regions (88%A–12%B, Supplemental Figs. S1C,J). Class 2 sites exhibit a faint boundary-like Hi-C contact pattern (Fig. 2C), weaker in magnitude compared to boundaries at cohesin occupancy sites in interphase, and similar to cohesin sites during meiosis (Supplemental Fig. S2F). These boundary-like patterns align with insulation score minima, and are more apparent with Class 2 binding sites derived from the B6 *Prdm9* allele (Supplemental Figs. S1B,C).

We also observe that particularly during meiosis, PRDM9 Class 2 sites – but not Class 1 – are enriched for overlaps with cohesin, CTCF and RNAPII ChIP-seq sites (Fig. 2D), each of which are themselves also biased toward A-compartment (Supplemental Fig. S3A). This enrichment is even more pronounced analyzing the B6 *Prdm9* allele (Supplemental Figs. S1D,E). Overlap with cohesin ChIP-seq sites potentially relates to the proposal that Class 2 binding represents axial tethering events[8], as cohesin is enriched at chromosomal axes during meiosis. However, this requires a cautious interpretation as we find that like Class 2 binding, cohesin ChIP-seq sites exhibit low cis/total ratio indicating high spatial accessibility. This is inconsistent with expectation for stable axial locations (Supplements S2). Unlike in yeast, our analysis suggests that mouse cohesin ChIP-seq site positions should not be interpreted as permanently axial loci throughout meiosis. Instead, loop positions during mammalian meiosis are likely highly stochastic between cells, lacking a set of reproducible axial positions[14].

Class 2 sites also exhibit strong enrichment for promoter and enhancer-associated histone marks such as H3K4me3/H3K4me1/H3K27me3, whereas enrichment is much more modest for PRDM9 Class 1 sites (Fig. 2E). Class 1 binding is expected to exhibit H3K4me3 enrichment due to PRDM9 methyltransferase activity at recombination hotspots[6, 7, 5]. While this is not observed in the CAST *Prdm9* allele due to genotype mismatch with chromHMM histone annotations derived from B6 mice, we did confirm H3K4me3 enrichment at B6 PRDM9 Class 1 sites (Supplemental Fig. S1M).

### DSB formation is enriched in active, spatially accessible chromosomal regions

We next investigated factors that affect the likelihood of PRDM9 Class 1 binding sites to generate DSBs during leptonema by partitioning PRDM9 Class 1 binding into quartiles with the highest and lowest respective DMC1 SSDS ChIP-seq scores (i.e., the most and least favoured PRDM9 sites for DSB formation) (Fig. 3A). Top quartile DSB-favoured sites are characterized by strong PRDM9 binding (Fig. 3A) and also exhibit stronger local shifts toward active, spatially accessible chromatin during zygonema relative to disfavoured sites (Figs. 3B,C). This further supports the idea that local active chromatin shifts observed during zygonema are related to PRDM9 methyltransferase activity, which is positively associated with DSB formation[9, 36]. In addition to local zygonema-specific effects, DSB-favoured sites appear to be biased at a more global scale towards active and spatially accessible chromosomal regions throughout interphase and meiosis (Fig. 3B).

**Figure 3:**
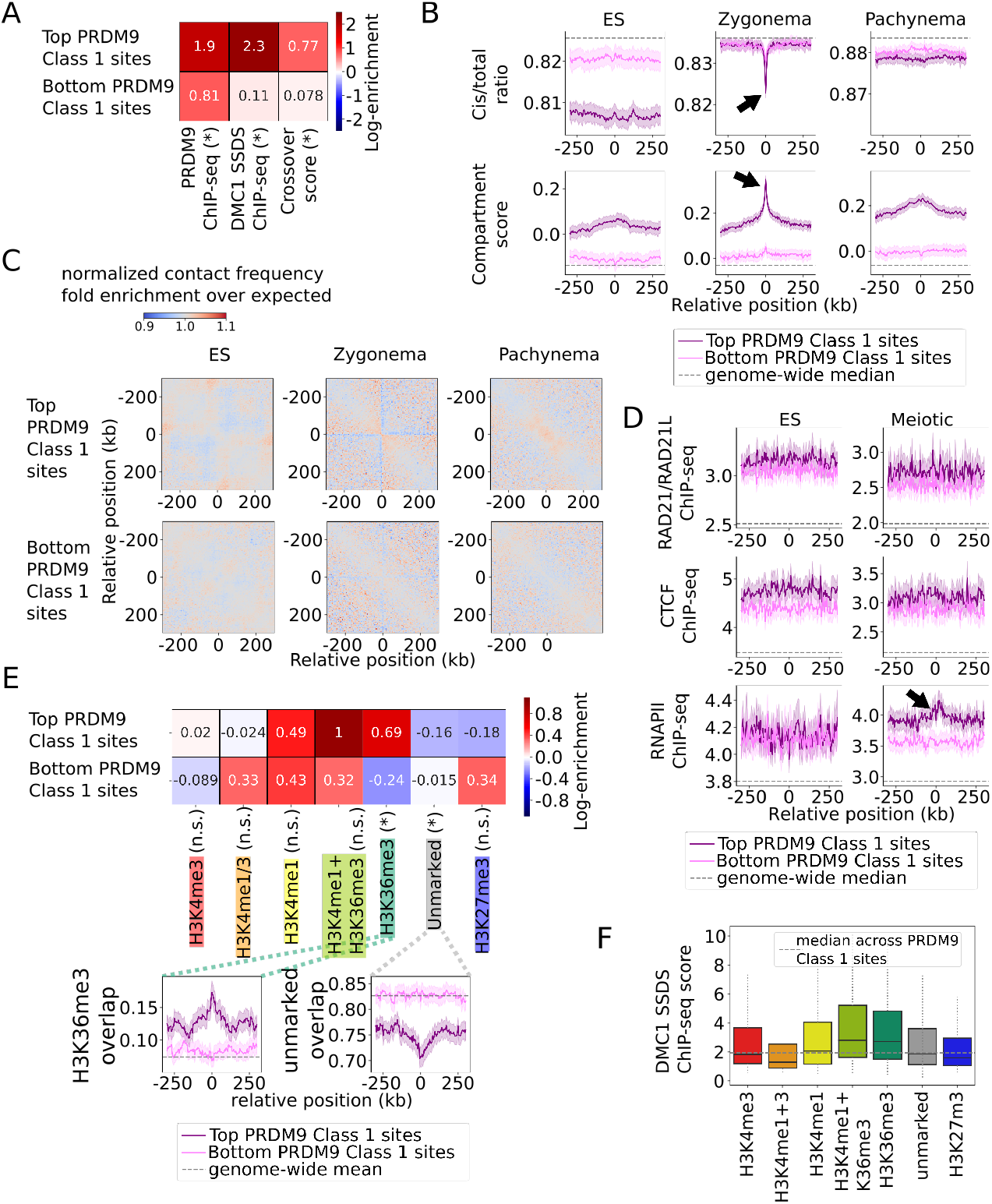
DSB formation is associated with active, spatially accessible chromatin. **A:** PRDM9 Class 1 sites are partitioned into the top and bottom quartiles of DMC1 SSDS ChIP-seq score, measuring DSB activity. Top (i.e., DSB-favoured) sites have more bound PRDM9 and greater likelihood of crossover formation. Heatmap shows log fold enrichment over 500 kb surrounding region, while (*) indicates Bonferroni-adjusted p<0.01 difference between top and bottom partitioned sites. **B:** Hi-C cis/total ratio (top) and compartment score (bottom), averaged across DSB-partitioned PRDM9 Class 1 sites, calculated for ES, zygonema and pachynema datasets. Shading represents 95% confidence intervals. Top DSB-favoured sites are associated with higher compartment score and lower cis/total ratio in general, black arrows indicate zygomena-specific shifts toward active spatially accessible chromatin, enhanced at DSB-favoured sites. **C:** Normalized chromatin contact matrices, averaged across top (DSB-favoured) and bottom PRDM9 Class 1 sites, for embryonic stem cell (ES), zygonema and pachynema Hi-C datasets. **D:** Cohesin subunit (top), CTCF (middle), and RNAPII (bottom) ChIP-seq tracks, averaged across DSB-partitioned PRDM9 Class 1 sites, calculated for ES and meiotic datasets. Elevated RNAPII occupancy during meiosis appears to be associated with increased DSB formation (black arrow). **E:** Overlap of chromHMM histone annotations with DSB-partitioned PRDM9 Class 1 sites. Note DSB-favoured sites are depleted for unmarked chromatin, while enriched for H3K36me3 chromatin typical of gene bodies. Heatmap shows log fold enrichment over genome-wide mean, while (*) indicates Bonferroni-adjusted p<0.01 difference between most (top) and least (bottom) DSB-favoured sites. Insets plot overlap fraction averaged around DSB-partitioned PRDM9 Class 1 sites, shading represents 95% confidence intervals. **F:** Distribution of DMC1 ChIP-seq scores (i.e. DSB activity) at PRDM9 Class 1 sites split by chromHMM state. DSB formation is elevated within H3K36me3-marked chromatin characteristic of gene bodies compared to regions associated with promoters, enhancers, and unmarked DNA.

We find that higher meiotic RNAPII occupancy is associated with increased DSB activity, whereas CTCF and cohesin levels do not appear strongly associated (Fig. 3D). As a corollary of enrichment toward A-compartment, DSB-favoured sites appear closer on average to CTCF/cohesin/RNAPII ChIP-seq sites – all heavily A-compartment biased – than DSB-disfavoured sites in terms of genomic distance (Supplemental Fig. S3C). We find enrichment of H3K36me3 histone marks (characteristic of gene bodies) and depletion of unmarked chromatin at DSB-favoured sites (Fig. 3E). Among all chromHMM states, DSBs appear most favoured at hotspots associated with H3K36me3 (Fig. 3F). Collectively, these results support the view that DSBs are enriched in active/spatially accessible chromatin near gene-rich genomic regions.

### Crossovers are depleted in spatially accessible chromatin, especially gene bodies

We next explore how chromatin organization around DSBs formed during leptonema affects their likelihood of being selected as the site of crossover formation later during pachynema. By cross-referencing sites of DSB formation with genomic maps of crossovers, we partition DSB sites (total of 6044 bins) into quartiles based on their crossover likelihood (Fig. 4A). We find that the top quartile of crossover-favoured DSB sites are characterized by elevated cis/total ratio throughout meiosis and interphase, and modestly lower compartment scores (Fig. 4B). These sites also exhibit depleted Hi-C contacts within a broad +/−100kb window (Fig. 4C) particularly during pachynema. This indicates that DSBs in chromosomal regions which adopt on average less spatially accessible configurations are favoured for crossover formation, in stark contrast with the clear enrichment of DSBs at PRDM9 binding sites in active, spatially accessible chromatin.

**Figure 4:**
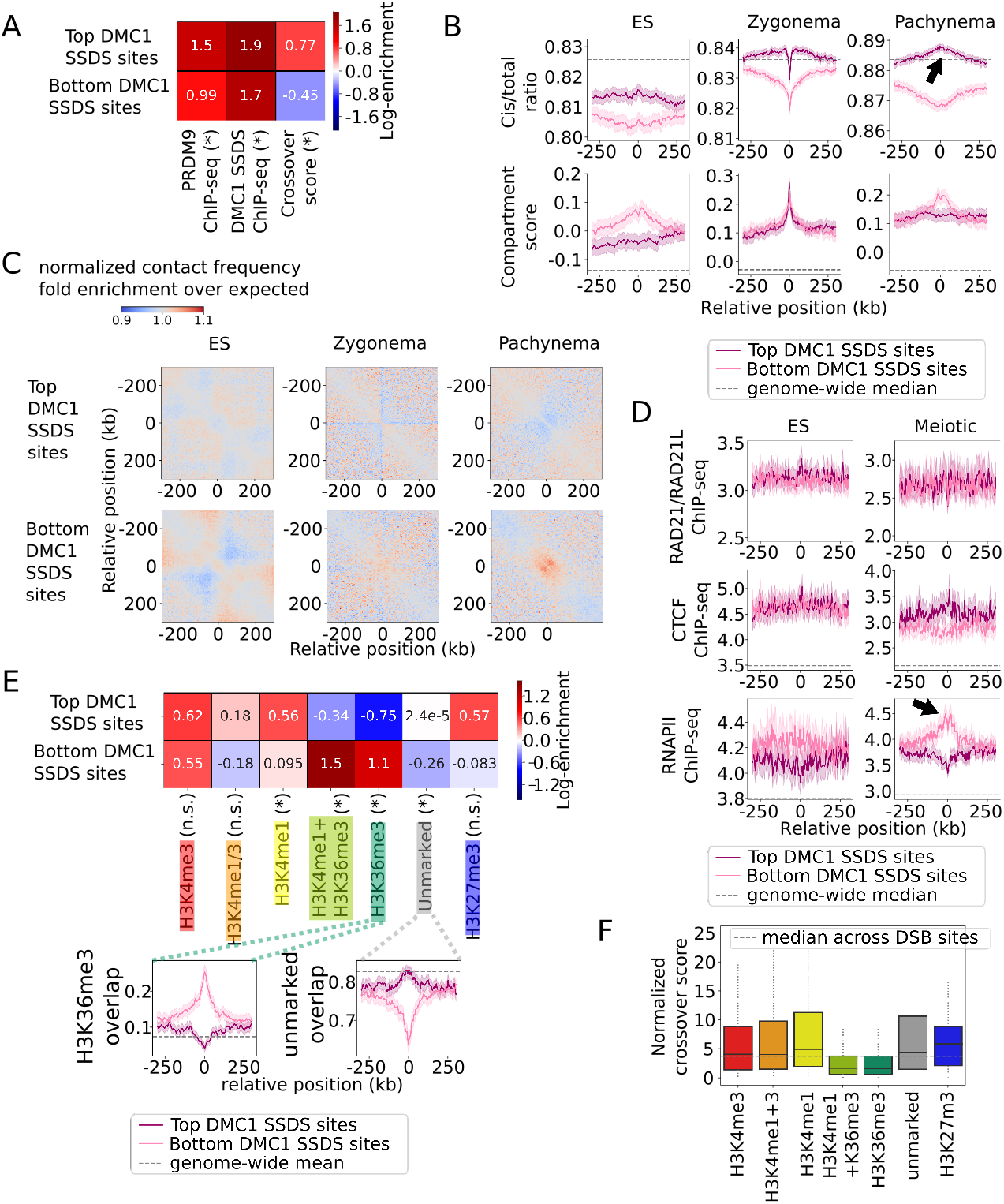
Spatially accessible chromatin is depleted for crossover formation, particularly at gene bodies. **A:** DSB sites (from DMC1 SSDS ChIP-seq) are partitioned into the top and bottom quartiles for crossover likelihood. Top (i.e., crossover-favoured) sites have modestly stronger inherent DSB activity, and exhibit greater likelihood of crossover formation. Heatmap shows log fold enrichment over 500 kb surrounding region, while (*) indicates Bonferroni-adjusted p<0.01 difference between top and bottom partitioned sites. **B:** Hi-C cis/total ratio (top) and compartment score (bottom), averaged across crossover-partitioned DSB sites, calculated for ES, zygonema and pachynema datasets. Shading represents 95% confidence intervals. Top crossover-favoured sites are associated with higher cis/total ratio particularly during meiosis, indicative of reduced spatial accessibility (black arrow). **C:** Normalized chromatin contact matrices, averaged across top (crossover-favoured) and bottom DSB sites, for embryonic stem cell (ES), zygonema and pachynema Hi-C datasets. Note reduced contact frequency near top (crossover-favoured) loci, particularly during pachynema. **D:** Cohesin subunit (top), CTCF (middle), and RNAPII (bottom) ChIP-seq tracks, averaged across crossover-partitioned DSB sites, calculated for ES and meiotic datasets. Elevated RNAPII occupancy during meiosis appears to be associated with decreased crossover formation (black arrow). **E:** Overlap of chromHMM histone annotations with crossover-partitioned DSB sites. Note crossover-favoured sites are depleted for H3K36me3 chromatin typical of gene bodies, while relatively enriched for unmarked chromatin. Heatmap shows log fold enrichment over genome-wide mean, while (*) indicates Bonferroni-adjusted p<0.01 difference between most (top) and least (bottom) crossover-favoured sitess. Insets plot overlap fraction averaged around crossover-partitioned DSB sites, shading represents 95% confidence intervals. **F:** Distribution of crossover likelihood scores at DSB sites split by chromHMM state. Note clear depletion of crossover formation at H3K36me3 marked chromatin, characteristic of gene bodies. Analogous plot for DSB activity at PRDM9 Class 1 sites (Fig. 3F) shows that DSB formation is not depleted within H3K36me3-marked chromatin, confirming that depletion of recombination within gene bodies occurs at the crossover selection step rather than during DSB formation.

In further contrast with DSB formation, we find that higher meiotic RNAPII occupancy appears associated with *decreased* crossover activity (Fig. 4D). The same holds true for elevated overlap with H3K36me3 histone marks, characteristic of gene bodies (Fig. 4E). Depletion of crossover formation in H3K36me3 gene body regions is confirmed by comparing crossover scores at DSB sites across different chromHMM histone annotations, while validating that this gene-body recombination depletion is not occurring at the earlier stage of DSB formation (Fig. 4F).

Interestingly, DSB sites that co-localize with H3K36me3 histone marks are characterized by low cis/total ratio (i.e. high spatial accessibility), which is associated with reduced crossover formation. However, crossover depletion at low cis/total ratio loci is observed beyond H3K36me3 sites (Supplemental Fig. S4). Collectively, these results support the view that crossover formation is depleted in active and spatially accessible chromatin, particularly gene bodies, and that distinctly different chromatin environments favour DSB formation and crossover formation.

### Linear model with principal component analysis reveals recombination-associated chromatin features

To jointly assess the contributions of different chromatin structure variables to the likelihood of DSB formation at PRDM9 sites and crossover formation at DSB sites, we implemented a linear modeling approach that uses principal component analysis (PCA) and model selection. These techniques help us to address the problem that Hi-C scores (cis/total ratio, compartment score, insulation, FIRE), epigenetic states (chromHMM), and ChIP-seq signals (cohesin, CTCF, RNAPII) constitute a large compendium of chromatin features, many of which are highly correlated with each other (Supplemental Fig. S5A).

First, we apply PCA to extract the primary directions of variation in chromatin variables from both ES and meiotic time points. To emphasize variation across recombination hotspots (as opposed to genomewide variation), we applied PCA among the union of bins marked as either PRDM9 Class 1 or DMC1 SSDS ChIP-seq sites. We find that the first principal component (PC1), encompassing by far the highest explained variance (16.0%), is positively associated with many of the general trends of active and spatially accessible chromatin. Increased levels of PC1 correspond to lower cis/total ratio, higher compartment score, and increased active epigenetic marks (Fig. 5A). The second principal component (PC2, 3.7% explained variance) positively associates with the presence of gene bodies (H3K36me3), along with high Hi-C FIRE scores, and meiosis-specific decreases in cis/total ratio. The third principal component (PC3, 2.7% explained variance) reflects instances where cis/total ratio and compartment scores diverge from their PC1 association. Positive PC3 indicates when spatial accessibility is higher than expected given chromatin activity. Finally, the fourth principal component (PC4, 2.4% explained variance) appears to reflect local enrichment of meiotic cohesin, H3K4me3, and RNAPII, consistent with presence of active promoters.

**Figure 5:**
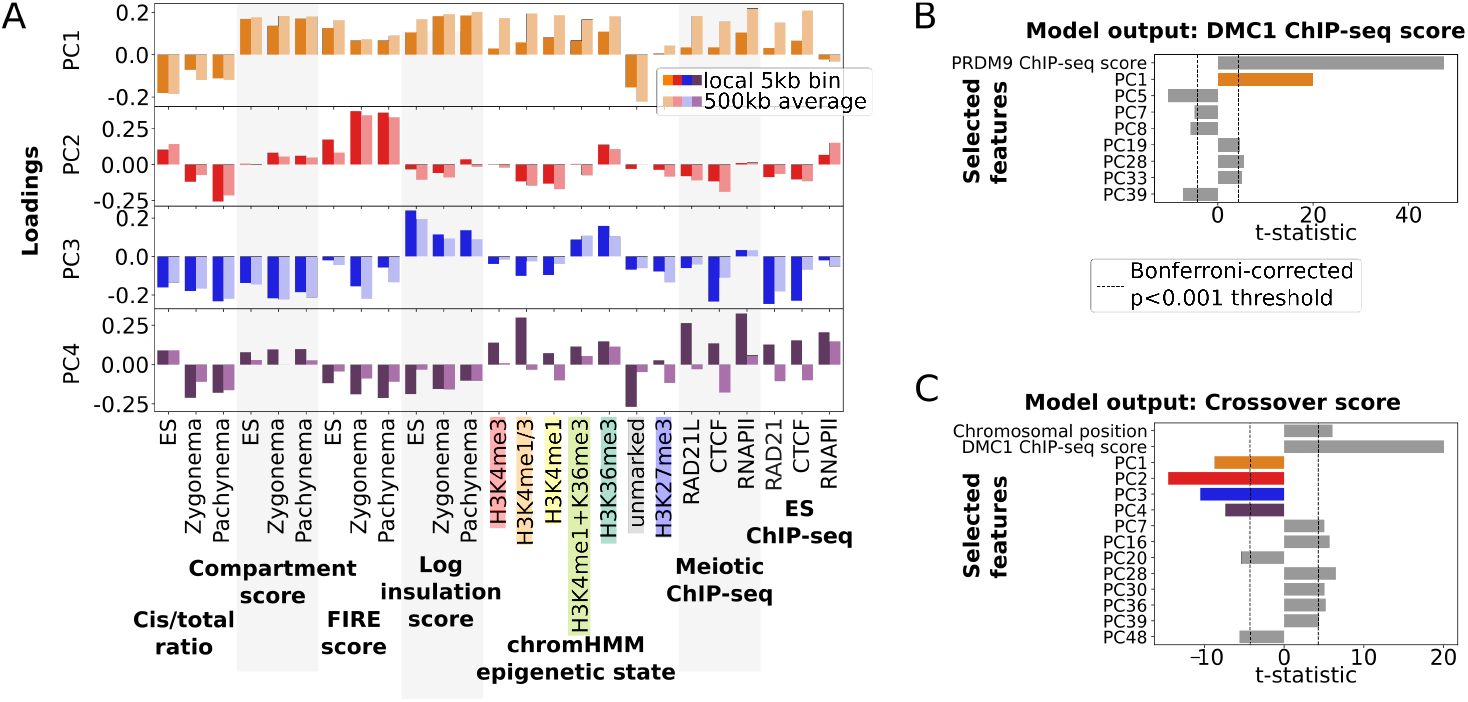
Principal component analysis with linear model reveals variable chromatin organization amongst recombination hotspots, and effect on recombination activity. **A:** Loadings for the top four principal components (PC1-PC4) of variation at recombination hotspots based on underlying chromatin organization variables (horizontal axis): Hi-C derived features (cis/total ratio, compartment score, FIRE score, and loginsulation score, for ES, zygonema and pachynema datasets), chromHMM epigenetic states, meiotic / ES ChIP-seq score for CTCF, RAD21(L) and RNAPII. Two values of each variable were included, one measuring local value at the recombination hotspot, the other a 500kb average around the hotspot. Positive PC1 loadings reflect presence of active chromatin, which is typically characterized by low cis/total ratio, high compartment, FIRE, and insulation scores, as well as increased histone modifications, cohesin and RNAPII. Positive PC2 loadings indicate presence of H3K36me3 histone marks typical of gene bodies, which tend to colocalize with increases in FIRE score, as well as meiotic specific decreases in cis/total ratio. PC3 reflects divergence from the typical correlation between activity and spatial accessibility – specifically, positive PC3 loadings indicate chromatin regions that are more spatially accessible (lower cis/total ratio) than expected given their activity (compartment score). Positive PC4 loadings indicate strong local enrichment of co-hesin/CTCF/RNAPII, characteristic of cohesin/CTCF/RNAPII occupancy sites. **B:** Results from a linear model for DSB activity (quantified by DMC1 SSDS ChIP-seq score) at PRDM9 Class 1 sites as a function of the principal components described in panel **A**, and adjusted for inherent PRDM9 binding variability (PRDM9 ChIP-seq score). Forward selection was used to choose statistically significant principal components to include in the model. Note that chromosomal position was not selected as significant, and the strongest predictors are positive inherent PRDM9 binding strength and positive PC1, reflecting increased DSB formation at hotspots in active, spatially accessible chromatin. PC colours as in **A**. **C:** Results from a linear model for crossover likelihood at DSB sites as a function of principal components and adjusted for centromeric proximity (chromosomal position) and inherent DSB variability (DMC1 SSDS ChIP-seq score). Forward selection was applied as in panel **B**. DMC1 and chromosomal position both positively predict crossover formation, reflective of inherent DSB variability and pericentromeric crossover depletion. PC1-4 all negatively predict crossover formation, reflecting crossover depletion at hotspots in active chromatin, particularly at gene bodies, promoters and spatially accessible regions. PC colours as in **A**.

After transforming chromatin variables into principal component space, we use a linear model with feature selection to identify statistically significant associations between principal components and either DSB or crossover formation. Applying this strategy to DSB formation at PRDM9 sites (Fig. 5B), we treat the 6421 PRDM9 Class 1 binding site bins as quasi-independent observations. We include PRDM9 binding score and chromosomal position in the model to adjust for inherent differences in PRDM9 binding and potential centromeric or telomeric proximity effects (mouse chromosomes are acrocentric, meaning chromosomal position can be used as a proxy for centromeric distance). We find that PRDM9 binding strength and PC1 are both strong predictors of DSB formation. The positive coefficient for PC1 confirms that DSBs are enriched in active chromatin. This effect is significant even after adjusting for differences in inherent PRDM9 binding strength at hotspots. Comparing explained variance between the naive model – only using PRDM9 ChIP-seq strength – and our selected model incorporating structural information, we find that *R^2^* increases from 27.1% to 35.4%.

Next, we apply the same approach to modeling the distribution of crossover likelihood at 6044 DSB sites (Fig. 5C), including DMC1 SSDS ChIP-seq score as a predictor to adjust for inherent differences in DSB formation. We find that PC1 (chromatin activity) exhibits a negative association with crossover likelihood, confirming that active and spatially accessible chromatin is disfavoured for crossovers, in contrast to its positive association with DSBs. Additionally, PC2-4 are also negative predictors of crossover likelihood, respectively indicating that gene bodies, promoters, and extra-spatially-accessible chromatin are disfavoured for crossover formation. We also find a strong positive correlation with chromosomal position as expected from crossover suppression near centromeres[37, 38]. Chromosomal position (along chromosomal arm) was only selected in the crossover prediction model (Fig. 5C), and not the DSB model (Fig. 5B), supporting earlier conclusions that pericentromeric regions are depleted for crossovers but not DSBs[37, 39, 40]. Comparing explained variance between a naive model – only using DMC1 ChIP-seq strength and chromosomal position – and our selected model incorporating structural information, we find that *R*^2^ increases from 9.1% to 19.6%.

### Organization of A and B compartment genomic regions exhibit different chromosomal loop lengths

Given the importance of chromatin activity and spatial accessibility in recombination, as well as the somewhat puzzling low cis/total ratio observed at meiotic cohesin peaks, we investigated further how A and B compartment regions are organized in the context of brush-loop meiotic chromosomes. While the density and physical size of loops appear relatively consistent along the axes of meiotic chromosomes[41], previous cytological analysis suggests that active gene-rich chromatin regions are over-represented along the axis (i.e., stretched relative to inactive gene-poor chromatin)[42, 43]. By analyzing contact frequency versus distance curves[44] in meiotic Hi-C data, we find the maxima of the derivatives occurs on average ~3-fold shorter in the A-compartment compared with the B-compartment. This suggests that meiotic loops in the A-compartment may be shorter in terms of their average lengths in base pairs (Supplemental Fig. S6). Assuming loops have consistent spacing and physical size along the axis[41] irrespective of their compartment, this would be consistent with reported over-representation and relative decondensation of active chromatin along the physical length of chromosomal axes[42, 43]. Enrichment of axial loci in active chromatin can also help resolve the otherwise puzzling low cis/total ratio at cohesin ChIP-seq sites, given that active, decondensed chromatin is generally associated with lower cis/total ratio[45, 27].

## Discussion

We test the hypothesis that features of 3D genome organization are associated with meiotic recombination in the mammalian genome. Our analyses reveal distinct associations with PRDM9 binding, DSB formation, and crossover formation. We confirm these observations with multiple regression, and analyze contact frequency decay to help situate these events relative to the meiotic brush-loop structure (Fig. 6A). This work complements earlier multi-factorial analysis of DSB and crossover formation at hotspots[12, 13] by demonstrating that in addition to local chromatin environment, features of genome folding beyond the immediate vicinity of hotspots (e.g., > 5kb) are significantly associated with differences in recombination.

**Figure 6:**
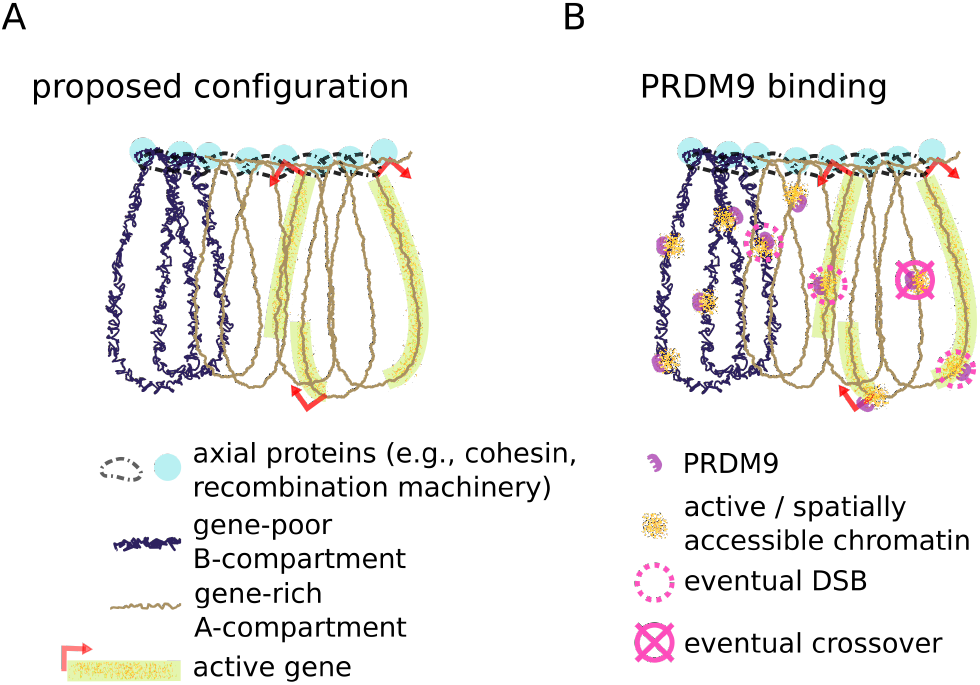
Proposed framework relating mammalian meiotic chromosomal architecture and recombination. **A:** In early leptonema, meiotic chromosomes adopt a brush-loop architecture, with cohesin and recombination machinery located at the axis. Loops in the gene-rich A-compartment have on average fewer base-pairs than B-compartment. Accordingly, A and B compartment regions depicted here represent roughly equal genomic lengths despite greater number of A-compartment loops. Physical size of A and B compartment loops may remain comparable due to relaxed linear packing density in gene-rich A-compartment. **B:** Concurrently during leptonema, PRDM9 binds to recombination hotspots (Class 1 Binding) across both A and B-compartment regions. Binding causes local increases in chromatin activity and spatial accessibility. Schematic depicts hypothetical example of ten total binding events across A and B compartment regions. After PRDM9 binding, a subset of binding loci are recruited to DSB machinery at the axis and form DSBs. This subset is biased toward gene-rich A-compartment. Later during pachynema, a single crossover point is selected from the DSBs formed earlier, avoiding DSBs formed in gene-body regions.

PRDM9 Class 1 binding sites (recombination hotspots) occur with similar frequencies in both A and B compartments. We find that hotspots are also locally associated with a transient shift toward increased compartment scores and reduced cis/total ratios during zygonema (Fig. 6B). This builds on previous observations of a simultaneous transient shift in patterns of histone trimethylation (e.g., H3K4me3) around PRDM9 binding sites in mammals[6, 7, 5, 36]. In contrast, PRDM9 Class 2 (non-hotspot) binding sites occur more frequently in the A-compartment, and often overlap with cohesin ChIP-seq sites. This may relate to the proposal that Class 2 binding sites correspond to axial tethering[8], though further work is required for clarification given the lack of reproducible axial positions in meiotic Hi-C[14].

We find DSB formation at hotspots is strongly associated with elevated spatial accessibility and confirm DSB enrichment in active (A-compartment) chromatin[14]. DSB formation is also favoured in hotspots with enhanced zygonema-specific shifts toward increased activity and spatial accessibility (Fig. 6B). We speculate that spatially accessible PRDM9 bound hotspots may be more likely to physically encounter DSB machinery.

In contrast to DSB formation, crossovers are favoured at loci with less spatially accessible chromatin with increased cis/total ratios on average. Spatially accessible gene body regions marked by H3K36me3 are particularly depleted for crossovers (Fig. 6B), consistent with earlier reports in plants[46]. Our results argue that crossover depletion at gene bodies occurs at the crossover selection stage rather than during DSB formation. This may be a general feature of 3D genome structure, as decreased Hi-C cis/total ratio is associated with crossover depletion among DSB sites beyond H3K36me3 regions as well.

Because crossovers reflect rare and contingent events, interpreting ensemble-average Hi-C datasets presents many caveats. In any given cell, only a fraction of PRDM9 binding sites are converted to DSBs, and out of these only a fraction are selected as crossovers. Therefore, the Hi-C signals we observe at DSB or crossover-favoured genomic coordinates are unlikely to directly reflect chromosomal configuration at individual DSB and crossover events, as even the most favoured coordinates are not sites of recombination in most cells. Additionally, we note that genomic positions of DSBs and cohesin ChIP sites appear to have relatively low cis/total ratios, despite the fact that immunofluorescent microscopy shows DSB machinery and cohesin subunits enriched along the axes [47, 48, 49]. In general, genomic loci with axial positions in a brush-loop structure would be expected to display high cis/total ratios characteristic of low spatial accessibility. This puzzling observation is in contrast with yeast meiosis, where cohesin ChIP sites display elevated cis/total ratios [50, 51], as expected for axially positioned loci in a loop-brush structure.

Our analyses of contact frequency curves in meiosis suggest that, on average, loops in active chromatin have fewer base-pairs than in inactive chromatin. We speculate that overall linear packing density of chromatin may be higher in B-compartment, compensating for base-pair length differences (Fig. 6A). This proposal is congruent with previously reported over-representation of A-compartment regions along chromosomal axes[42, 43], and aligns with the enrichment of meiotic cohesin peaks in A-compartment[17, 18]. Transcriptionally active genomic regions with decondensed chromatin are furthermore associated with increased spatial accessibility[45, 27]; together this may resolve the otherwise puzzling low cis/total ratio observed at meiotic cohesin ChIP-seq sites, which often overlap promoters. Nevertheless, this association between transcriptional activity and axial localization requires a cautious interpretation. Despite sharing many ChIP-seq sites with cohesin at promoters, RNAPII – which is additionally enriched in gene bodies – is broadly dispersed along chromosomal loops[52]. H3K4me3, which marks promoters and recombination hotspots, is present both at axial and loop positions[53], whereas H3K27me3, which typically associated with transcriptionally repressed promoters, localizes close to the axis[53]. Given these findings, we cannot currently rule out the possibility that cohesin is more uniformly distributed across active and inactive chromatin, yet is preferentially visible in transcriptionally active regions such as promoters due to hyper-ChIPability[54, 55].

We conclude by suggesting avenues for future experimental studies. First, modern microscopy methods such as DNA-paint and super-resolution FISH[56, 57, 58] would be useful to trace contiguous DNA regions and obtain direct evidence for differing numbers of base-pairs per loop in active and inactive regions. Second, our results thus far largely demonstrate associations between genome organization and recombination – perturbation experiments will be necessary to explore causality. For instance, to test the effects of chromatin activity and spatial accessibility, we envision experiments that target expression of genes near or overlapping recombination hotspots using CRISPR inhibition/activation tools[59, 60], in conjunction with chromosomal conformation capture and ChIP-seq experiments to observe downstream effects on meiotic genome folding, PRDM9 binding, DSB formation, and crossovers. Third, to explore the connection between PRDM9 methyltransferase activity and transient shifts in 3D genome organization, we imagine potential Hi-C and ChIP-seq experiments using heterozygous PRDM9 mutants with modified methyltransferase activity[61]. Alternatively, CRISPR-based epigenome modification strategies[62] can be repurposed to directly perturb histone marks around hotspots, exploring for example whether targeted local H3K4me3 deposition is sufficient to drive genome folding changes during meiosis to the extent observed in zygonema with PRDM9. Similar approaches could be used to investigate the effects of modified H3K36me3 levels for DSB and crossover formation. Finally, meiotic Hi-C for human chromosomes would enable an analysis of the potential interplay between *Prdm9* polymorphism [7] and clinically relevant dysregulation of 3D genome folding in meiosis.

## Methods

### Genome mapping and 5kb bins

All analyses were performed with the mouse mm10 genome assembly. `Cooler makebins` was used to generate a 5kb bin bed file corresponding to the mm10 genome. Using this bedfile as auxiliary input, `bedtools intersect` was used to convert unbinned genomic tracks to 5kb resolution, outputting the maximum score and coverage per bin for bedgraph and bed tracks respectively.

### ChIP-seq datasets

The encode chip-seq pipeline https://github.com/ENCODE-DCC/chip-seq-pipeline2 was used to process raw ChIP-seq reads into bigwig trace (fold change over input) and peak site files. Pre-processed peak bed and bigwig trace files were used where available, and lifted over to mm10 if needed. PRDM9 ChIP-seq sites were further classified as Class 1 and Class 2 by filtering for intersections with DMC1 or H3K4me3 binding as described previously[8].

### Generating crossover likelihood map from single-sperm-seq dataset

A 5kb resolution crossover likelihood map was generated using a list of mapped crossover loci from Yin *et al.*. These loci were derived using single sperm sequencing of B6xCAST hybrid mouse[35]. Crossover likelihood for each 5kb bin was determined by summing the number of intersecting crossovers, inversely weighted by the length of each crossover (i.e., giving prominence to sharply localized crossovers).

### chromHMM chromatin epigenetic states

Chromatin state was obtained from Mouse ENCODE project, using the segmentation outputs from the testis-specific 7-state chromHMM model, along with ES-derived for interphase reference[32].

### Hi-C analysis

Raw Hi-C reads from ES[24] and meiosis (zygonema, pachynema)[14, 17] were converted using the `Hi-CUP` pipeline into `cooler` format at 5kb resolution and analyzed using cooltools.

### Compartment vector calling using fine grain eigenvectors

While compartment vectors for ES Hi-C samples were successfully generated using `cooltools eigdecomp` for whole chromosomes (i.e., traditional eigenvector decomposition), the approach failed to generate reasonable compartment vectors for several chromosomes in the pachynema and zygonema dataset due to the dropoff in signal of the Hi-C matrices beyond 10Mb contact distances. Therefore, we implemented the “fine grain eigenvector” approach presented by Wang. et al.[16] which calculates eigenvectors using 10Mb x 10Mb chunks along the Hi-C contact map.

### Visualization and plotting

`Seaborn` and `matplotlib` were used to generate meta-averaged Hi-C matrices and genomic tracks. `PyGenomeTracks` was used generate example genome tracks at particular loci. Bins with minimal read counts are filtered out by `cooler` during Hi-C matrix balancing and ignored during visualization.

### PCA and linear model with feature selection

Values for all chromatin organization variables (see Figure 5A, horizontal axis) were calculated for recombination hotspot bins, defined as the union of PRDM9 Class 1 and DMC1 SSDS ChIP-seq sites. Variables were pre-processed with standard scaling (0-mean, 1-standard deviation), followed by principal component analysis using the `sklearn.decomposition` package to extract loadings for each principal component. Principal components were calculated for all PRDM9 Class 1 sites and supplied (alongside PRDM9 ChIP-seq score and chromosomal position) as predictors to a linear model predicting DMC1 SSDS ChIP-seq score using forward variable selection. Briefly, the forward selection process begins with a null model and adds variables one-by-one by choosing the most statistically significant predictor at each step into an updated least squares model, implemented using `statsmodels.OLS`. The selection process converges when none of the remaining predictors passes the statistical significance threshold, in this case p<0.001 with Bonferroni correction (significance threshold = 0.001/n, for *n* variables). The selected predictors and their respective t-statistics are then reported. The process was repeated for DMC1 SSDS ChIP-seq sites in a model which used principal components alongside DMC1 SSDS ChIP-seq score and chromosomal position as predictor variables to a linear model predicting crossover likelihood score.

## Availability of data and materials

All datasets pertaining to this manuscript are previously published and described in Supplemental Information S0.

## Author’s contributions

XJ, GF and KP contributed to the design and implementation of the research, to the analysis of the results and to the writing of the manuscript.

## Acknowledgements

We thank S. Whalen, L. Chumpitaz-Diaz, K. Keough, S. Schalbetter and M. Neale for helpful discussions regarding data analysis and feedback on the manuscript.

## References

[1] Lam, I., Keeney, S.: Mechanism and Regulation of Meiotic Recombination Initiation. Cold Spring Harbor Perspectives in Biology 7(1) (2015). doi:10.1101/cshperspect.a016634

[2] Potapova, T., Gorbsky, G.J.: The Consequences of Chromosome Segregation Errors in Mitosis and Meiosis. Biology 6(1), 12 (2017). doi:10.3390/biology6010012

[3] Petronis, A.: Alzheimer’s disease and down syndrome: from meiosis to dementia. Experimental neurology 158(2), 403–413 (1999). doi:10.1006/exnr.1999.7128

[4] Baudat, F., Imai, Y., de Massy, B.: Meiotic recombination in mammals: localization and regulation. Nature Reviews Genetics 14, 794 (2013)

[5] Parvanov, E.D., Petkov, P.M., Paigen, K.: PRMD9 Controls Activation of Mammalian Recombination Hotspots. Science 327(5967), 835–835 (2010). doi:10.1126/science.1181495

[6] Myers, S., Bowden, R., Tumian, A., Bontrop, R.E., Freeman, C., MacFie, T.S., McVean, G., Donnelly, P.: Drive Against Hotspot Motifs in Primates Implicates the PRDM9 Gene in Meiotic Recombination. Science 327(5967), 876–879 (2010). doi:10.1126/science.1182363

[7] Baudat, F., Buard, J., Grey, C., Fledel-Alon, A., Ober, C., Przeworski, M., Coop, G., de Massy, B.: PRDM9 is a major determinant of meiotic recombination hotspots in humans and mice. Science(New York, N.Y.) 327(5967), 836–840 (2010). doi:10.1126/science.1183439

[8] Grey, C., Clement, J.A.J., Buard, J., Leblanc, B., Gut, I., Gut, M., Duret, L., de Massy, B.: In vivo binding of PRDM9 reveals interactions with noncanonical genomic sites. Genome research 27(4), 580–590 (2017). doi:10.1101/gr.217240.116

[9] Diagouraga, B., Clément, J.A.J., Duret, L., Kadlec, J., de Massy, B., Baudat, F.: PRDM9 Methyltransferase Activity Is Essential for Meiotic DNA Double-Strand Break Formation at Its Binding Sites. Molecular Cell 69(5), 853–8656 (2018). doi:10.1016/j.molcel.2018.01.033

[10] Baudat, F., de Massy, B.: Regulating double-stranded DNA break repair towards crossover or noncrossover during mammalian meiosis. Chromosome Research 15(5), 565–577 (2007). doi:10.1007/s10577-007-1140-3

[11] Li, R., Bitoun, E., Altemose, N., Davies, R.W., Davies, B., Myers, S.R.: A high-resolution map of non-crossover events reveals impacts of genetic diversity on mammalian meiotic recombination. Nature Communications 10(1), 3900 (2019). doi:10.1038/s41467-019-11675-y

[12] Yamada, S., Kim, S., Tischfield, S.E., Jasin, M., Lange, J., Keeney, S.: Genomic and chromatin features shaping meiotic double-strand break formation and repair in mice. Cell cycle (Georgetown, Tex.) 16(20), 1870–1884 (2017). doi:10.1080/15384101.2017.1361065

[13] Hinch, A.G., Zhang, G., Becker, P.W., Moralli, D., Hinch, R., Davies, B., Bowden, R., Donnelly, P.: Factors influencing meiotic recombination revealed by whole-genome sequencing of single sperm. Science 363(6433), 8861 (2019). doi:10.1126/science.aau8861

[14] Patel, L., Kang, R., Rosenberg, S.C., Qiu, Y., Raviram, R., Chee, S., Hu, R., Ren, B., Cole, F., Corbett, K.D.: Dynamic reorganization of the genome shapes the recombination landscape in meiotic prophase. Nature Structural and Molecular Biology 26(3), 164–174 (2019). doi:10.1038/s41594-019-0187-0

[15] Alavattam, K.G., Maezawa, S., Sakashita, A., Khoury, H., Barski, A., Kaplan, N., Namekawa, S.H.: Attenuated chromatin compartmentalization in meiosis and its maturation in sperm development. Nature Structural and Molecular Biology 26(3), 175–184 (2019). doi:10.1038/s41594-019-0189-y

[16] Wang, Y., Wang, H., Zhang, Y., Du, Z., Si, W., Fan, S., Qin, D., Wang, M., Duan, Y., Li, L., Jiao, Y., Li, Y., Wang, Q., Shi, Q., Wu, X., Xie, W.: Reprogramming of Meiotic Chromatin Architecture during Spermatogenesis. Molecular Cell 73(3), 547–5616 (2019). doi:10.1016/j.molcel.2018.11.019

[17] Vara, C., Paytuví-Gallart, A., Cuartero, Y., Le Dily, F., Garcia, F., Salva-Castro, J., Gomez-H, L., Julia, E., Moutinho, C., Aiese Cigliano, R., Sanseverino, W., Fornas, O., Pendás, A.M., Heyn, H., Waters, P.D., Marti-Renom, M.A., Ruiz-Herrera, A.: Three-Dimensional Genomic Structure and Cohesin Occupancy Correlate with Transcriptional Activity during Spermatogenesis. Cell reports 28(2), 352-3679 (2019). doi:10.1016/j.celrep.2019.06.037

[18] Luo, Z., Wang, X., Jiang, H., Wang, R., Chen, J., Chen, Y., Xu, Q., Cao, J., Gong, X., Wu, J., Yang, Y., Li, W., Han, C., Cheng, C.Y., Rosenfeld, M.G., Sun, F., Song, X.: Reorganized 3D Genome Structures Support Transcriptional Regulation in Mouse Spermatogenesis. iScience 23(4), 101034 (2020). doi:10.1016/j.isci.2020.101034

[19] Møens, P.B., Pearlman, R.E.: Chromatin organization at meiosis. BioEssays 9(5), 151–153 (1988). doi:10.1002/bies.950090503

[20] Blat, Y., Protacio, R.U., Hunter, N., Kleckner, N.: Physical and Functional Interactions among Basic Chromosome Organizational Features Govern Early Steps of Meiotic Chiasma Formation. Cell 111, 791–802 (2002)

[21] Tock, A.J., Henderson, I.R.: Hotspots for Initiation of Meiotic Recombination. Frontiers in genetics 9(November), 521 (2018). doi:10.3389/fgene.2018.00521

[22] Grey, C., Baudat, F., de Massy, B.: PRDM9, a driver of the genetic map. PLOSGenetics 14(8), 1007479 (2018)

[23] Slotman, J.A., Paul, M.W., Carofiglio, F., de Gruiter, H.M., Vergroesen, T., Koornneef, L., van Cappellen, W.A., Houtsmuller, A.B., Baarends, W.M.: Super-resolution imaging of RAD51 and DMC1 in DNA repair foci reveals dynamic distribution patterns in meiotic prophase. PLOSGenetics 16(6), 1008595 (2020)

[24] Bonev, B., Mendelson Cohen, N., Szabo, Q., Fritsch, L., Papadopoulos, G.L., Lubling, Y., Xu, X., Lv, X., Hugnot, J.-P., Tanay, A., Cavalli, G.: Multiscale 3D Genome Rewiring during Mouse Neural Development. Cell 171(3), 557–57224 (2017). doi:10.1016/j.cell.2017.09.043

[25] Nitzsche, A., Paszkowski-Rogacz, M., Matarese, F., Janssen-Megens, E.M., Hubner, N.C., Schulz, H., de Vries, I., Ding, L., Huebner, N., Mann, M., Stunnenberg, H.G., Buchholz, F.: RAD21 Cooperates with Pluripotency Transcription Factors in the Maintenance of Embryonic Stem Cell Identity. PLOSONE 6(5), 19470 (2011)

[26] Shen, Y., Yue, F., McCleary, D.F., Ye, Z., Edsall, L., Kuan, S., Wagner, U., Dixon, J., Lee, L., Lobanenkov, V.V., Ren, B.: A map of the cis-regulatory sequences in the mouse genome. Nature 488(7409), 116–120 (2012). doi:10.1038/nature11243

[27] Kalhor, R., Tjong, H., Jayathilaka, N., Alber, F., Chen, L.: Genome architectures revealed by tethered chromosome conformation capture and population-based modeling. Nature biotechnology 30(1), 90–98 (2011). doi:10.1038/nbt.2057

[28] Falk, M., Feodorova, Y., Naumova, N., Imakaev, M., Lajoie, B.R., Leonhardt, H., Joffe, B., Dekker, J., Fudenberg, G., Solovei, I., Mirny, L.A.: Heterochromatin drives compartmentalization of inverted and conventional nuclei. Nature 570(7761), 395–399 (2019). doi:10.1038/s41586-019-1275-3

[29] Lieberman-Aiden, E., Van Berkum, N.L., Williams, L., Imakaev, M., Ragoczy, T., Telling, A., Amit, I., Lajoie, B.R., Sabo, P.J., Dorschner, M.O., Sandstrom, R., Bernstein, B., Bender, M.A., Groudine, M., Gnirke, A., Stamatoyannopoulos, J., Mirny, L.A., Lander, E.S., Dekker, J., Miiro, G., Serwanga, J., Pozniak, A., Aacphee, D., Manigart, L., Aawananyanda, E., Karita, A., Inwoley, W., Jaoko, Dehovitz, L.G., Bekker, P., Pitisuttithum, R., Paris, S.A., Sabo, P.J.: Comprehensive Mapping of Long-Range Interactions Reveals Folding Principles of the Human Genome. Science 326(5950), 289–293 (2009)

[30] Crane, E., Bian, Q., McCord, R.P., Lajoie, B.R., Wheeler, B.S., Ralston, E.J., Uzawa, S., Dekker, J., Meyer, B.J.: Condensin-driven remodelling of X chromosome topology during dosage compensation. Nature 523(7559), 240–244 (2015). doi:10.1038/nature14450

[31] Schmitt, A.D., Hu, M., Jung, I., Xu, Z., Qiu, Y., Tan, C.L., Li, Y., Lin, S., Lin, Y., Barr, C.L., Ren, B.: A Compendium of Chromatin Contact Maps Reveals Spatially Active Regions in the Human Genome. Cellreports 17(8), 2042–2059 (2016). doi:10.1016/j.celrep.2016.10.061

[32] Yue, F., Cheng, Y., Breschi, A., Vierstra, J., Wu, W., Ryba, T., Sandstrom, R., Ma, Z., Davis, C., Pope, B.D., Shen, Y., Pervouchine, D.D., Djebali, S., Thurman, R.E., Kaul, R., Rynes, E., Kirilusha, A., Marinov, G.K., Williams, B.A., Trout, D., Amrhein, H., Fisher-Aylor, K., Antoshechkin, I., DeSalvo, G., See, L.-H., Fastuca, M., Drenkow, J., Zaleski, C., Dobin, A., Prieto, P., Lagarde, J., Bussotti, G., Tanzer, A., Denas, O., Li, K., Bender, M.A., Zhang, M., Byron, R., Groudine, M.T., McCleary, D., Pham, L., Ye, Z., Kuan, S., Edsall, L., Wu, Y.-C., Rasmussen, M.D., Bansal, M.S., Kellis, M., Keller, C. A., Morrissey, C.S., Mishra, T., Jain, D., Dogan, N., Harris, R.S., Cayting, P., Kawli, T., Boyle, A.P., Euskirchen, G., Kundaje, A., Lin, S., Lin, Y., Jansen, C., Malladi, V.S., Cline, M.S., Erickson, D. T., Kirkup, V.M., Learned, K., Sloan, C.A., Rosenbloom, K.R., Lacerda de Sousa, B., Beal, K., Pignatelli, M., Flicek, P., Lian, J., Kahveci, T., Lee, D., James Kent, W., Ramalho Santos, M., Herrero, J., Notredame, C., Johnson, A., Vong, S., Lee, K., Bates, D., Neri, F., Diegel, M., Canfield, T., Sabo, P.J., Wilken, M.S., Reh, T.A., Giste, E., Shafer, A., Kutyavin, T., Haugen, E., Dunn, D., Reynolds, A.P., Neph, S., Humbert, R., Scott Hansen, R., De Bruijn, M., Selleri, L., Rudensky, A., Josefowicz, S., Samstein, R., Eichler, E.E., Orkin, S.H., Levasseur, D., Papayannopoulou, T., Chang, K.-H., Skoultchi, A., Gosh, S., Disteche, C., Treuting, P., Wang, Y., Weiss, M.J., Blobel, G.A., Cao, X., Zhong, S., Wang, T., Good, P.J., Lowdon, R.F., Adams, L.B., Zhou, X.-Q., Pazin, M.J., Feingold, E.A., Wold, B., Taylor, J., Mortazavi, A., Weissman, S.M., Stamatoyannopoulos, J.A., Snyder, M.P., Guigo, R., Gingeras, T.R., Gilbert, D.M., Hardison, R.C., Beer, M.A., Ren, B., Consortium, T.M.E.: A comparative encyclopedia of DNA elements in the mouse genome. Nature 515(7527), 355–364 (2014). doi:10.1038/nature13992

[33] Margolin, G., Khil, P.P., Kim, J., Bellani, M.A., Camerini-Otero, R.D.: Integrated transcriptome analysis of mouse spermatogenesis. BMC genomics 15, 39 (2014). doi:10.1186/1471-2164-15-39

[34] Smagulova, F., Brick, K., Pu, Y., Camerini-Otero, R.D., Petukhova, G.V.: The evolutionary turnover of recombination hot spots contributes to speciation in mice. Genes& development 30(3), 266–280 (2016). doi:10.1101/gad.270009.115

[35] Yin, Y., Jiang, Y., Lam, K.-W.G., Berletch, J.B., Disteche, C.M., Noble, W.S., Steemers, F.J., Camerini-Otero, R.D., Adey, A.C., Shendure, J.: High-Throughput Single-Cell Sequencing with Linear Amplification. Molecular Cell 76(4), 676–69010 (2019). doi:10.1016/j.molcel.2019.08.002

[36] Chen, Y., Lyu, R., Rong, B., Zheng, Y., Lin, Z., Dai, R., Zhang, X., Xie, N., Wang, S., Tang, F., Lan, F., Tong, M.-H.: Refined spatial temporal epigenomic profiling reveals intrinsic connection between PRDM9-mediated H3K4me3 and the fate of double-stranded breaks. Cell Research 30(3), 256–268 (2020). doi:10.1038/s41422-020-0281-1

[37] Blitzblau, H.G., Bell, G.W., Rodriguez, J., Bell, S.P., Hochwagen, A.: Mapping of Meiotic SingleStranded DNA Reveals Double-Strand-Break Hotspots near Centromeres and Telomeres. Current Biology 17(23), 2003–2012 (2007). doi:10.1016/j.cub.2007.10.066

[38] Nambiar, M., Smith, G.R.: Repression of harmful meiotic recombination in centromeric regions. Seminarsin Cell& Developmental Biology 54, 188–197 (2016). doi:10.1016/j.semcdb.2016.01.042

[39] Talbert, P.B., Henikoff, S.: Centromeres Convert but Don’t Cross. PLOS Biology 8(3), 1000326 (2010)

[40] Vincenten, N., Kuhl, L.-M., Lam, I., Oke, A., Kerr, A.R.W., Hochwagen, A., Fung, J., Keeney, S., Vader, G., Marston, A.L.: The kinetochore prevents centromere-proximal crossover recombination during meiosis. eLife 4, 10850 (2015). doi:10.7554/eLife.10850

[41] Zickler, D., Kleckner, N.: Meiotic Chromosomes: Integrating Structure and Function. Annual Review of Genetics 33(1), 603–754 (1999). doi:10.1146/annurev.genet.33.1.603

[42] Luciani, J.M., Guichaoua, M.R., Cau, P., Devictor, B., Salagnon, N.: Differential elongation of autosomal pachytene bivalents related to their DNA content in human spermatocytes. Chromosoma 97(1), 19–25 (1988). doi:10.1007/BF00331791

[43] Fransz, P.F., Armstrong, S., de Jong, J.H., Parnell, L.D., van Drunen, C., Dean, C., Zabel, P., Bisseling, T., Jones, G.H.: Integrated Cytogenetic Map of Chromosome Arm 4S of A.thaliana: Structural Organization of Heterochromatic Knob and Centromere Region. Cell 100(3), 367–376 (2000). doi:10.1016/S0092-8674(00)80672-8

[44] Gassler, J., Brandão, H.B., Imakaev, M., Flyamer, I.M., Ladstätter, S., Bickmore, W.A., Peters, J.-M., Mirny, L.A., Tachibana, K.: A mechanism of cohesin-dependent loop extrusion organizes zygotic genome architecture. TheEMBO journal 36(24), 3600–3618 (2017). doi:10.15252/embj.201798083

[45] Mahy, N.L., Perry, P.E., Bickmore, W.A.: Gene density and transcription influence the localization of chromatin outside of chromosome territories detectable by FISH. Journal of Cell Biology 159(5), 753–763 (2002). doi:10.1083/jcb.200207115

[46] Wijnker, E., Velikkakam James, G., Ding, J., Becker, F., Klasen, J.R., Rawat, V., Rowan, B.A., de Jong, D.F., de Snoo, C.B., Zapata, L., Huettel, B., de Jong, H., Ossowski, S., Weigel, D., Koornneef, M., Keurentjes, J.J.B., Schneeberger, K.: The genomic landscape of meiotic crossovers and gene conversions in Arabidopsis thaliana. eLife 2, 01426 (2013). doi:10.7554/eLife.01426

[47] Lee, J., Hirano, T.: RAD21L, a novel cohesin subunit implicated in linking homologous chromosomes in mammalian meiosis. Journal of Cell Biology 192(2), 263–276 (2011). doi:10.1083/jcb.201008005

[48] Ishiguro, K.-i., Kim, J., Fujiyama-Nakamura, S., Kato, S., Watanabe, Y.: A new meiosis-specific cohesin complex implicated in the cohesin code for homologous pairing. EMBO reports 12(3), 267–275 (2011). doi:10.1038/embor.2011.2

[49] Hinch, A.G., Becker, P.W., Li, T., Moralli, D., Zhang, G., Bycroft, C., Green, C., Keeney, S., Shi, Q., Davies, B., Donnelly, P.: The Configuration of RPA, RAD51, and DMC1 Binding in Meiosis Reveals the Nature of Critical Recombination Intermediates. Molecular Cell 79(4), 689–70110 (2020). doi:10.1016/j.molcel.2020.06.015

[50] Muller, H., Scolari, V.F., Agier, N., Piazza, A., Thierry, A., Mercy, G., Descorps-Declere, S., Lazar-Stefanita, L., Espeli, O., Llorente, B., Fischer, G., Mozziconacci, J., Koszul, R.: Characterizing meiotic chromosomes’ structure and pairing using a designer sequence optimized for Hi-C. Molecular systems biology 14(7), 8293–8293 (2018). doi:10.15252/msb.20188293

[51] Schalbetter, S.A., Füdenberg, G., Baxter, J., Pollard, K.S., Neale, M.J.: Principles of meiotic chromosome assembly revealed in S. cerevisiae. Nature Communications 10(1), 4795 (2019). doi:10.1038/s41467-019-12629-0

[52] van der Laan, R., Uringa, E.J., Wassenaar, E., Hoogerbrugge, J.W., Sleddens, E., Odijk, H., Roest, H.P., de Boer, P., Hoeijmakers, J.H.J., Grootegoed, J.A., Baarends, W.M.: Ubiquitin ligase Rad18Sc localizes to the XY body and to other chromosomal regions that are unpaired and transcriptionally silenced during male meiotic prophase. Journal of Cell Science 117(21), 5023–5033 (2004). doi:10.1242/jcs.01368

[53] Prakash, K., Fournier, D., Redl, S., Best, G., Borsos, M., Tiwari, V.K., Tachibana-Konwalski, K., Ketting, R.F., Parekh, S.H., Cremer, C., Birk, U.J.: Superresolution imaging reveals structurally distinct periodic patterns of chromatin along pachytene chromosomes. Proceedings of the National Academy of Sciences 112(47), 14635–14640 (2015). doi:10.1073/pnas.1516928112

[54] Teytelman, L., Thurtle, D.M., Rine, J., van Oudenaarden, A.: Highly expressed loci are vulnerable to misleading ChIP localization of multiple unrelated proteins. Proceedings of the National Academy of Sciences 110(46), 18602–18607 (2013). doi:10.1073/pnas.1316064110

[55] Jain, D., Baldi, S., Zabel, A., Straub, T., Becker, P.B.: Active promoters give rise to false positive ‘Phantom Peaks’ in ChIP-seq experiments. Nucleic acids research 43(14), 6959–6968 (2015). doi:10.1093/nar/gkv637

[56] Schnitzbauer, J., Strauss, M.T., Schlichthaerle, T., Schueder, F., Jungmann, R.: Super-resolution microscopy with DNA-PAINT. NatureProtocols 12(6), 1198–1228 (2017). doi:10.1038/nprot.2017.024

[57] Beliveau, B.J., Boettiger, A.N., Avendaño, M.S., Jungmann, R., McCole, R.B., Joyce, E.F., Kim-Kiselak, C., Bantignies, F., Fonseka, C.Y., Erceg, J., Hannan, M.A., Hoang, H.G., Colognori, D., Lee, J.T., Shih, W.M., Yin, P., Zhuang, X., Wu, C.-t.: Single-molecule super-resolution imaging of chromosomes and in situ haplotype visualization using Oligopaint FISH probes. NatureCommunications 6(1), 7147 (2015). doi:10.1038/ncomms8147

[58] Albert, P.S., Zhang, T., Semrau, K., Rouillard, J.-M., Kao, Y.-H., Wang, C.-J.R., Danilova, T.V., Jiang, J., Birchler, J.A.: Whole-chromosome paints in maize reveal rearrangements, nuclear domains, and chromosomal relationships. Proceedings of the National Academy of Sciences 116(5), 1679–1685 (2019). doi:10.1073/pnas.1813957116

[59] Larson, M.H., Gilbert, L.A., Wang, X., Lim, W.A., Weissman, J.S., Qi, L.S.: CRISPR interference (CRISPRi) for sequence-specific control of gene expression. Nature Protocols 8(11), 2180–2196 (2013). doi:10.1038/nprot.2013.132

[60] Maeder, M.L., Linder, S.J., Cascio, V.M., Fu, Y., Ho, Q.H., Joung, J.K.: CRISPR RNA–guided activation of endogenous human genes. Nature Methods 10(10), 977–979 (2013). doi:10.1038/nmeth.2598

[61] Thibault-Sennett, S., Yu, Q., Smagulova, F., Cloutier, J., Brick, K., Camerini-Otero, R.D., Petukhova, G.V.: Interrogating the Functions of PRDM9 Domains in Meiosis. Genetics 209(2), 475–487 (2018). doi:10.1534/genetics.118.300565

[62] Hilton, I.B., D’Ippolito, A.M., Vockley, C.M., Thakore, P.I., Crawford, G.E., Reddy, T.E., Gersbach, C.A.: Epigenome editing by a CRISPR-Cas9-based acetyltransferase activates genes from promoters and enhancers. Nature Biotechnology 33(5), 510–517 (2015). doi:10.1038/nbt.3199

