## Supplementary Information for "Genome-wide variability in recombination activity is associated with meiotic chromatin organization"

#### S0 Datasets used in integrative analysis

The table below lists all datasets used in our integrative analysis. In choosing datasets to focus on when similar instances were available (e.g., meiotic Hi-C[1, 2, 3]), we seek where possible to maintain consistency in terms of mouse strain background. For instance, the PRDM9, DMC1 (DSB) and crossover datasets all analyzed recombination in a B6 x CAST hybrid context.

| Data type | Cell/tissue type | Mouse strain information | Citation | Notes |
|---|---|---|---|---|
| Hi-C | E14TG2a Embryonic stem (ES) cell line | 129/Ola background | Bonev et al.[4] | Used for ES Hi-C in Figs. 2-4. |
| Hi-C | Zygotene and pachytene spermatocytes | B6 x CAST | Patel et al.[1] | Used for zygonema, pachynema Hi-C in Figs. 1-4, mouse strain background matches PRDM9 and DSB datasets.[5, 6] |
| Hi-C | Pachytene spermatocytes | B6 x PWK | Wang et al.[2] | Used for additional pachynema Hi-C in Supplements S2. |
| Hi-C | Lepto/zygotene, pachy/diplotene spermatocytes and round spermatids | B6 | Vara et al.[3] | Used in Supplements S2. |

| | | | | |
|---|---|---|---|---|
| Hi-C | Round spermatids | B6 | Alavattam et al.[7] | Used in Supplements S2, processed to 20kb instead of 5kb resolution due to 6-cutter (rather than 4-cutter) restriction enzyme. |
| ChIP-seq (anti-CTCF, anti-REC8, anti-RAD21L) | Pachy/diplotene spermatocytes | B6 | Vara et al.[3] | Used for meiotic ChIP-seq in Figs. 1-4. |
| ChIP-seq (anti-RNAPII) | Testis extract from 16 dpp mice | B6 | Margolin et al.[8] | Used for meiotic ChIP-seq in Figs. 1-4. |
| ChIP-seq (anti-CTCF, RAD21) | ES-R1 ES cell line | 129 background | Nitzsche et al.[9] | Used for ES ChIP-seq in Figs. 2-4. |
| ChIP-seq (anti-RNAPII) | Bruce4 ES cell line | B6 background | Shen et al.[10] | Used for ES ChIP-seq in Figs. 2-4. |
| ChIP-seq (anti-PRDM9) | Testis extract from 13 dpp mice | B6 and RJ2 – RJ2 expresses PRMD9-CAST on C57BL/10 background | Grey et al.[5] | RJ2 data used for PRDM9-CAST binding signal in Figs. 1-3, B6 data analyzed for PRDM9-B6 in Supplements S1. |
| SSDS ChIP-seq (anti-DMC1) | Testis extract adult mice | B6 x CAST | Smagulova et al.[6] | Data used for double-strand-break (DSB) signal in Figs. 1-4. |
| Single-cell whole genome sequencing | Mature 1C sperm | B6 x CAST | Yin et al.[11] | Data used for crossover signal in Figs. 1-4. |

### S1 Comparing B6 and CAST *Prdm9* allele results

While most of the PRDM9-related results presented in the main text are from ChIP-seq of the CAST *Prdm9* allele on a B6 background (RJ2 strain[5]), we also analyze patterns for the native B6 allele on a B6 genetic background. Overall, similar chromatin organization is observed around binding sites of the B6 *Prdm9* allele compared to CAST – here we discuss some differences between the two alleles. As reported in the original publication[5], fewer peaks are detected with the native B6 allele - 1881 Class 1, 718 Class 2 - than the CAST allele - 7218 Class 1, 2651 Class 2. Note mm9-mm10 lift-over process resulted in slight differences in counts from original publication, and that number of genomic sites (i.e., bins with peaks) is by definition less than the total number of peaks. A similar B6 genotype serves as the background for both the B6 and CAST *Prdm9* strains on which PRDM9 ChIP-seq was performed[5], and it is believed that reduced B6 binding may be related to potential hotspot erosion (i.e., loss over evolutionary timescales of B6 PRDM9 protein binding sites on its native background) and reported dominance of CAST *Prdm9* allele over B6[5, 6]. The increased number of CAST sites relative B6 is a key reason why we focus on the CAST results to augment signal in our observations averaged across sites.

In addition to fewer overall binding sites, B6 PRDM9 Class 1 sites tend to be less likely to form DSBs and crossovers as compared to CAST (Fig. S1A), consistent with previously reported dominance of the CAST allele. B6 PRDM9 Class 1 binding sites exhibit overall very similar signals to CAST counterpart in terms of 3D organization measured by Hi-C (Figs. S1B,C). Compared with CAST allele results, B6 Class 1 sites also exhibit local zygonema-specific shifts toward spatially accessible active configuration of low cis/total ratio and high compartment score. B6 Class 2 sites exhibit even stronger enrichment toward active, spatially accessible chromatin compared to CAST Class 2. B6 Class 2 sites also exhibit stronger boundary-like contact patterns than CAST Class 2 (Fig. S1B), accordingly with more distinct insulation score minima (Fig. S1C). With Class 2 binding sites, B6 PRDM9 protein binding appears even more enriched for / localized near CTCF/cohesin/RNAPII sites as compared with CAST (Figs. S1D,E), with B6 Class 2 sites often overlapping cohesin, CTCF and RNAPII sites. In accordance with our results indicating spatially accessible chromatin at RAD21L sites (Supplements S2), we find that B6 PRDM9 Class 2 sites are associated with increased spatial accessibility (lower cis/total ratio), as compared with their CAST counterparts (Fig. S1C).

Additionally, B6 PRDM9 Class 1 sites exhibit a much stronger enrichment towards H3K4me3 histone marks as determined using the chromHMM annotations from mouse testis tissue[12] (Figs. S1F,G). Typically, such H3K4me3 annotations are characteristic of promoter regions, however in this particular case the B6-specific enrichment suggests the signal is due to H3K4me3 methylation during meiosis caused by PRDM9 binding. This is because the mouse genotype used to produce chromHMM histone annotations was B6, rather than CAST, and as a result would only register H3K4me3 recruited at B6 PRDM9 Class 1 binding sites. In line with this reasoning, we do not see this strong overlap between B6 PRDM9 Class 1 sites and H3K4me3 chromHMM regions when using an interphase (ES) chromHMM annotation set (Fig. S1H), as PRDM9 binding (and associated H3K4me3 recruitment) does not occur during interphase. Furthermore, by splitting H3K4me3 chromHMM annotated regions into ES-specific, testis-specific, and shared, we find that the vast majority of B6 PRDM9 Class 1 sites are found in testis-specific H3K4me3 regions (Fig. S1I) consistent with these regions representing PRDM9-driven H3K4me3 trimethylation during meiosis. We also do not observe a strong enrichment toward A-compartment with B6 PRMD9 Class 1 binding as compared to CAST (Fig. S1J), further indicating the elevated H3K4me3 signal for B6 is not due to active-promoter enrichment.

In addition to our analysis with CAST PRDM9 sites, we also partition B6 PRDM9 Class 1 binding sites into the most and least DSB favoured quartiles. This analysis for B6 (Figs. S1K-M) reveals very similar trends with those from the CAST allele presented in the main text. Increased PRDM9 binding, enrichment of active A-compartment, and zygonema specific remodeling are all observed in the DSB-favoured B6 PRDM9 Class 1 sites. Consistent with our earlier results related to testis-specific H3K4me3, we find that specific to B6, DSB-favoured PRDM9 Class 1 sites are enriched for H3K4me3. Such sites are also depleted for unmarked quiescent chromatin similar to CAST.

#### S2 Chromosomal organization around cohesin occupancy sites

Cohesin is involved in the assembly of chromosomal axes during meiosis, and localizes at the axial core of the brush-loop structure during meiosis[13, 14, 3, 15, 16]. We analyze the ChIP-seq data for the meiosis-specific cohesin subunit RAD21L in pachytene-diplotene spermatocytes[3], as well as of its interphase counterpart RAD21 in ES cells[9]. We also considered REC8, another meiotic-specific cohesin kleisin subunit. Earlier microscopy results indicate that spatial occupancy patterns of RAD21L and REC8 are mutually exclusive on individual chromosomes[15, 16]. Our analysis based on bulk ChIP-seq data, however, indicates that at 5kb bin resolution, REC8 and RAD21L sites often overlap (Fig. S2A), consistent with original analysis of the dataset [3]. Follow-up work could explore whether there exists a universal occupancy pattern dictating mutual exclusion of REC8 versus RAD21L occupancy, or whether cell-to-cell differences in occupancy patterns dominate. By comparison, meiotic RAD21L and REC8 cohesin sites overlap much less frequently with interphase RAD21 sites (Fig. S2A).

We find that meiosis-specific cohesin sites exhibit low cis/total ratios (Fig. S2B). This is puzzling, because we expect that sites with preferred axial positioning would display elevated cis/total ratios reflective of decreased spatial accessibility. This observation also contrasts with those for yeast meiosis, where Rec8 cohesin sites are associated with local maxima in the cis/total ratio [17].

Digging deeper, the differences in average spatial accessibility across cohesin types during pachynema[1] appear related to the enrichment of RAD21L sites at active promoter-like regions marked by H3K4me3 (Fig. S2C) and away from unmarked (i.e., generally more quiescent) chromatin regions. Splitting the cohesin sites by chromHMM group prior to averaging, we observe that cis/total ratio trends are more similar across different cohesin subunits, in particular the meiotic subunits REC8 and RAD21L (Fig. S2D). Indeed, elevated cis/total ratio patterns are observed at RAD21L sites not associated with H3K4me3 or H3K36me3 marks typical of active promoters and gene bodies (Fig. S2D), much like interphase RAD21. This suggests that decreased cis/total ratio at meiotic RAD21L sites may be associated with RAD21L's increased colocalization with active promoter/enhancer-like histone marks and transcriptional activity relative to interphase RAD21. RAD21L sites also exhibit a modest enrichment for H3K27me3 marks (Fig. S2C) typical of repressed promoter regions, suggesting that enrichment of meiotic cohesin towards promoters may occur irrespective of active-vs.-repressed promoter state. H3K27me3-marked RAD21L sites are characterized by increased cis/total ratio (Fig. S2D).

Other explanations for low meiotic cis/total at RAD21L sites can include technical aspects of dataset collection. In particular, the RAD21L ChIP-seq dataset was generated from pachytene/diplotene spermatocytes[3], representing a later stage of meiotic prophase compared to the zygonema and pachynema Hi-C datasets from Patel et al.[1]. Therefore, it is possible that stable cohesin accumulation at RAD21L ChIP-seq sites is present in later stage pachytene/diplotene spermatocytes, but not in zygonema or early pachynema. Indeed, previous work suggests that ongoing loop extrusion occurs during earlier stages of meiotic prophase I[1]. Consistent with loop extrusion continuing until late prophase, we detect locally elevated cis/total ratio at RAD21L sites in post-meiotic round spermatids, pachytene/diplotene stage spermatocytes, as compared with low cis/total ratio in early prophase (Fig. S2E). We find a similar local elevation in another recently published pachytene Hi-C dataset published by Wang et al.[2] (Fig. S2B). We speculate this dataset may capture spermatocytes at a slightly later stage of pachynema.

Motivated by Hi-C patterns around cohesin sites during interphase, we next test for evidence of boundary demarcation at meiotic cohesin sites and enriched contacts between site pairs by considering Hi-C maps averaged around these features. We observe both boundary demarcation at individual cohesin sites during meiosis, as reported by Vara *et al.*[3], as well as enriched Hi-C contacts between proximal cohesin site pairs for RAD21, RAD21L and REC8 sites (Fig. S2F). These signals are more evident in pachynema than zygonema, especially for meiosis-specific subunit RAD21L, though overall much weaker than analogous RAD21-based signals in ES cells (i.e., interphase). Similar patterns are also observed in the pachytene Hi-C dataset by Wang et al. (Fig. S2F)[2]. Note in the main text we focus on the Patel *et al.* meiotic Hi-C contact maps due to strain matching with recombination datasets (B6xCAST). Given the REC8/RAD21L site overlap, averaged Hi-C signals at REC8 sites are similar to, albeit slightly weaker than, RAD21L (Fig. S2F).

Since previous work reported a relationship between cohesin positioning and transcriptional direction in yeast[18], we next explored whether similar trends could be observed in mammalian meiosis. By averaging meiotic Hi-C maps at RAD21L sites that uniquely intersect either positive or negative strand transcription start sites (TSS), we find that boundary signals occur both upstream and downstream regardless of strandedness, though downstream Hi-C contact frequency is modestly enriched for RAD21L sites intersecting +strand TSS and vice versa (Fig. S2G). This signal may be due to presence of enriched Hi-C contacts near gene bodies downstream of TSS. No clear differences are observed between convergent versus divergent RAD21L site pairs, which both show enrichment (Fig. S2G) – thus we are unable to clearly conclude a link between transcriptional direction and cohesin occupancy.

In addition to weak boundaries in Hi-C contact maps, we also observe minima in the log-insulation score (Fig. S2B) at RAD21L sites, as expected given the boundary signals described above. Insulation minima are observed in both meiotic and interphase (ES) Hi-C. This opens the possibility that these loci are partially predetermined by cohesin occupancy prior to meiotic entry, though sample contamination by different cell types or meiotic stages cannot be ruled out. Meanwhile, ES RAD21 cohesin site loci exhibit very strong boundary demarcation and dot-loop-contact signals in matching ES Hi-C, but a much weaker signal in meiotic Hi-C (Fig. S2F).

Overall, these observations indicate that pachytene/diplotene stage cohesin ChIP-seq sites, particularly for meiosis-specific RAD21L, *should not* be directly interpreted as stable locations of cohesin occupancy – and by extension the brush-loop axis – throughout meiosis. Cis/total ratios at cohesin sites exhibit minima during early prophase, and appear to increase during later prophase as well as in post-meiotic Hi-C (Figs. S2B,E). This suggests a potential configuration where these promoter and CTCF-enriched genomic positions increasingly arrive at the cohesin-anchored axis during meiosis, perhaps due to an increasing tendency for these loci to stall cohesin extrusion. Noting that the chromatin boundary and loop-enrichment signals at RAD21L sites / site pairs are more apparent during pachynema compared to zygonema (Fig. S2F), we speculate that these ChIP-seq sites may be characterized by increasing cohesin accumulation – and by extension axial localization – as meiotic prophase I progresses.

#### S3 Cohesin, CTCF and RNAPII peaks enriched in A-compartment

Along with RAD21, REC8 and RAD21L cohesin subunits, meiotic CTCF and RNAPII occupancy sites are heavily biased toward A-compartment chromatin (Fig. S3A). An interesting corollary of A-compartment cohesin enrichment is that other A-compartment enriched features are on average more likely to be close to a cohesin site. For instance, we demonstrate that in terms of genomic distance, PRDM9 Class 2 binding sites are on average closer to cohesin/CTCF/RNAPII sites than Class 1 (Fig. S3B). Similarly, DSB-favoured PRDM9 Class 1 binding sites are on average closer to cohesin/CTCF/RNAPII/Class 2 sites than DSB-disfavoured (Fig. S3C).

#### S4 Cis/total ratio of DSB sites split by chromHMM state

In the main text, we discuss how selection of crossovers from DSBs is depleted in genomic regions with low cis/total ratio (i.e., high spatial accessibility) particularly as measured from meiotic Hi-C, as well as in regions with H3K36me3 histone annotation from the mouse testis chromHMM dataset. Here we explore the relationship between these two observations - specifically, we seek to address whether DSB sites in H3K36me3 chromatin are inherently characterized by low cis/total ratio, and if so whether cis/total ratio is a relevant factor for crossover selection in chromHMM annotated regions beyond H3K36me3 regions. To investigate this, we first split DSB sites by their chromHMM annotation thus forming seven groups. For each individual group, we partitioned by crossover likelihood score, and plot the averaged pachynema cis/total ratio for the top and bottom quartile DSB sites of each group (Fig. S4). We note that H3K36me3 associated DSB sites are indeed characterized by minima in cis/total ratio. However, we also note that the tendency of low cis/total ratio sites to be depleted for crossover selection is observed in all chromHMM groups, not just H3K36me3-associated. Thus it appears that high spatial accessibility of chromatin during meiosis is associated with depleted crossover formation in general.

#### S5 Collinearity between variables describing chromatin organization and principal component analysis

A total of 50 variables (see x-axis of Fig. 5A) describe chromatin organization at each hotspot site, which are defined as the union of PRDM9 Class 1 sites (for both CAST and B6) and DMC1 sites (representing DSBs). To deal with collinearity between these variables (Fig. S5A), PCA is performed across hotspot sites, transforming these variables into a 50-dimensional principal component space. We also plot explained variances (Fig. S5B) of each principal component.

#### S6 Differences in loop lengths between A and B-compartment

Earlier cytological observations of meiotic chromosomes indicated two findings: (i) loop densities are relatively constant along chromosomal axes[19], and (ii) gene-rich chromatin – roughly equating to A-compartment – is over-represented (i.e., stretched) along the physical length of the axis[20, 21]. This suggests that for the same genomic distance, A-compartment chromatin forms more loops than B-compartment chromatin, implying that loops in A-compartment regions should have fewer base-pairs than B-compartment. To explore whether such a configuration can be observed in meiotic Hi-C datasets, we apply the $P(s)$ derivative method for estimating loop lengths[22, 1]. Briefly, by calculating the derivative of the contact frequency versus distance curve – commonly referred to as $P(s)$ curve – the genomic distance where the derivative equals zero can be used as an estimate of loop length.

We apply this technique to meiotic Hi-C contact matrices masking A-compartment and B-compartment regions respectively, and find indeed that there exists an approximately 2-3 fold difference in the number of base pairs in A versus B-compartment loops in zygonema (Fig. S6A). This finding is replicated in multiple pachynema Hi-C datasets (Figs. S6B-D), and is consistent with the bias in cohesin sites toward A-compartment (Fig. S3A).

#### References


[1] Lucas Patel, Rhea Kang, Scott C. Rosenberg, Yunjiang Qiu, Ramya Raviram, Sora Chee, Rong Hu, Bing Ren, Francesca Cole, and Kevin D. Corbett. Dynamic reorganization of the genome shapes the recombination landscape in meiotic prophase. *Nature Structural and Molecular Biology*, 26(3):164–174, 2019.

[2] Yao Wang, Hanben Wang, Yu Zhang, Zhenhai Du, Wei Si, Suixing Fan, Dongdong Qin, Mei Wang, Yanchao Duan, Lufan Li, Yuying Jiao, Yuanyuan Li, Qiujun Wang, Qinghua Shi, Xin Wu, and Wei Xie. Reprogramming of Meiotic Chromatin Architecture during Spermatogenesis. *Molecular Cell*, 73(3):547–561.e6, 2019.

[3] Covadonga Vara, Andreu Paytuví-Gallart, Yasmina Cuartero, François Le Dily, Francisca Garcia, Judit Salvà-Castro, Laura Gómez-H, Eva Julià, Catia Moutinho, Riccardo Aiese Cigliano, Walter Sanseverino, Oscar Fornas, Alberto M Pendás, Holger Heyn, Paul D Waters, Marc A Marti-Renom, and Aurora Ruiz-Herrera. Three-Dimensional Genomic Structure and Cohesin Occupancy Correlate with Transcriptional Activity during Spermatogenesis. *Cell reports*, 28(2):352–367.e9, jul 2019.

[4] Boyan Bonev, Netta Mendelson Cohen, Quentin Szabo, Lauriane Fritsch, Giorgio L Papadopoulos, Yaniv Lubling, Xiaole Xu, Xiaodan Lv, Jean-Philippe Hugnot, Amos Tanay, and Giacomo Cavalli. Multiscale 3D Genome Rewiring during Mouse Neural Development. *Cell*, 171(3):557–572.e24, oct 2017.

[5] Corinne Grey, Julie A J Clément, Jérôme Buard, Benjamin Leblanc, Ivo Gut, Marta Gut, Laurent Duret, and Bernard de Massy. In vivo binding of PRDM9 reveals interactions with noncanonical genomic sites. *Genome research*, 27(4):580–590, apr 2017.

[6] Fatima Smagulova, Kevin Brick, Yongmei Pu, R. Daniel Camerini-Otero, and Galina V. Petukhova. The evolutionary turnover of recombination hot spots contributes to speciation in mice. *Genes & development*, 30(3):266–280, feb 2016.

[7] Kris G. Alavattam, So Maezawa, Akihiko Sakashita, Haia Khoury, Artem Barski, Noam Kaplan, and Satoshi H. Namekawa. Attenuated chromatin compartmentalization in meiosis and its maturation in sperm development. *Nature Structural and Molecular Biology*, 26(3):175–184, 2019.

[8] Gennady Margolin, Pavel P Khil, Joongbaek Kim, Marina A Bellani, and R Daniel Camerini-Otero. Integrated transcriptome analysis of mouse spermatogenesis. *BMC genomics*, 15:39, jan 2014.

[9] Anja Nitzsche, Maciej Paszkowski-Rogacz, Filomena Matarese, Eva M Janssen-Megens, Nina C Hubner, Herbert Schulz, Ingrid de Vries, Li Ding, Norbert Huebner, Matthias Mann, Hendrik G Stunnenberg, and Frank Buchholz. RAD21 Cooperates with Pluripotency Transcription Factors in the Maintenance of Embryonic Stem Cell Identity. *PLOS ONE*, 6(5):e19470, may 2011.

[10] Yin Shen, Feng Yue, David F McCleary, Zhen Ye, Lee Edsall, Samantha Kuan, Ulrich Wagner, Jesse Dixon, Leonard Lee, Victor V Lobanenkov, and Bing Ren. A map of the cis-regulatory sequences in the mouse genome. *Nature*, 488(7409):116–120, 2012.

[11] Yi Yin, Yue Jiang, Kwan-Wood Gabriel Lam, Joel B Berletch, Christine M Disteche, William S Noble, Frank J Steemers, R Daniel Camerini-Otero, Andrew C Adey, and Jay Shendure. High-Throughput Single-Cell Sequencing with Linear Amplification. *Molecular Cell*, 76(4):676–690.e10, nov 2019.

[12] Feng Yue, Yong Cheng, Alessandra Breschi, Jeff Vierstra, Weisheng Wu, Tyrone Ryba, Richard Sandstrom, Zhihai Ma, Carrie Davis, Benjamin D Pope, Yin Shen, Dmitri D Pervouchine, Sarah Djebali, Robert E Thurman, Rajinder Kaul, Eric Rynes, Anthony Kirilusha, Georgi K Marinov, Brian A Williams, Diane Trout, Henry Amrhein, Katherine Fisher-Aylor, Igor Antoshechkin, Gilberto DeSalvo, Lei-Hoon See, Meagan Fastuca, Jorg Drenkow, Chris Zaleski, Alex Dobin, Pablo Prieto, Julien Lagarde, Giovanni Bussotti, Andrea Tanzer, Olgert Denas, Kanwei Li, M A Bender, Miaohua Zhang, Rachel Byron, Mark T Groudine, David McCleary, Long Pham, Zhen Ye, Samantha Kuan, Lee Edsall, Yi-Chieh Wu, Matthew D Rasmussen, Mukul S Bansal, Manolis Kellis, Cheryl A Keller, Christapher S Morrissey, Tejaswini Mishra, Deepti Jain, Nergiz Dogan, Robert S Harris, Philip Cayting, Trupti Kawli, Alan P Boyle, Ghia Euskirchen, Anshul Kundaje, Shin Lin, Yiing Lin, Camden Jansen, Venkat S Malladi, Melissa S Cline, Drew T Erickson, Vanessa M Kirkup, Katrina Learned, Cricket A Sloan, Kate R Rosenbloom, Beatriz Lacerda de Sousa, Kathryn Beal, Miguel Pignatelli, Paul Flicek, Jin Lian, Tamer Kahveci, Dongwon Lee, W James Kent, Miguel Ramalho Santos, Javier Herrero, Cedric Notredame, Audra Johnson, Shinny Vong, Kristen Lee, Daniel Bates, Fidencio Neri, Morgan Diegel, Theresa Canfield, Peter J Sabo, Matthew S Wilken, Thomas A Reh, Erika Giste, Anthony Shafer, Tanya Kutyavin, Eric Haugen, Douglas Dunn, Alex P Reynolds, Shane Neph, Richard Humbert, R Scott Hansen, Marella De Bruijn, Licia Selleri, Alexander Rudensky, Steven Josefowicz, Robert Samstein, Evan E Eichler, Stuart H Orkin, Dana Levasseur, Thalia Papayannopoulou, Kai-Hsin Chang, Arthur Skoultchi, Srikanta Gosh, Christine Disteche, Piper Treuting, Yanli Wang, Mitchell J Weiss, Gerd A Blobel, Xiaoyi Cao, Sheng Zhong, Ting Wang, Peter J Good, Rebecca F Lowdon, Leslie B Adams, Xiao-Qiao Zhou, Michael J Pazin, Elise A Feingold, Barbara Wold, James Taylor, Ali Mortazavi, Sherman M Weissman, John A Stamatoyannopoulos, Michael P Snyder, Roderic Guigo, Thomas R Gingeras, David M Gilbert, Ross C Hardison, Michael A Beer, Bing Ren, and The Mouse ENCODE Consortium. A comparative encyclopedia of DNA elements in the mouse genome. *Nature*, 515(7527):355–364, 2014.

[13] Elena Llano, Yurema Herrán, Ignacio García-Tuñón, Cristina Gutiérrez-Caballero, Enrique de Álava, José Luis Barbero, John Schimenti, Dirk G de Rooij, Manuel Sánchez-Martín, and Alberto M Pendás. Meiotic cohesin complexes are essential for the formation of the axial element in mice. *The Journal of cell biology*, 197(7):877–885, jun 2012.

[14] J Suja and J Barbero. Cohesin Complexes and Sister Chromatid Cohesion in Mammalian Meiosis. In *Genome Dynamics*, volume 5, pages 94–116. 2009.

[15] Jibak Lee and Tatsuya Hirano. RAD21L, a novel cohesin subunit implicated in linking homologous chromosomes in mammalian meiosis. *Journal of Cell Biology*, 192(2):263–276, jan 2011.

[16] Kei-ichiro Ishiguro, Jihye Kim, Sally Fujiyama-Nakamura, Shigeaki Kato, and Yoshinori Watanabe. A new meiosis-specific cohesin complex implicated in the cohesin code for homologous pairing. *EMBO reports*, 12(3):267–275, mar 2011.

[17] Stephanie A Schalbetter, Geoffrey Fudenberg, Jonathan Baxter, Katherine S Pollard, and Matthew J Neale. Principles of meiotic chromosome assembly revealed in S. cerevisiae. *Nature Communications*, 10(1):4795, 2019.

[18] Xiaoji Sun, Lingzhi Huang, Tovah E Markowitz, Hannah G Blitzblau, Doris Chen, Franz Klein, and Andreas Hochwagen. Transcription dynamically patterns the meiotic chromosome-axis interface. *eLife*, 4:e07424, aug 2015.

[19] D Zickler and N Kleckner. Meiotic Chromosomes: Integrating Structure and Function. *Annual Review of Genetics*, 33(1):603–754, dec 1999.

[20] J M Luciani, M R Guichaoua, P Cau, B Devictor, and N Salagnon. Differential elongation of autosomal pachytene bivalents related to their DNA content in human spermatocytes. *Chromosoma*, 97(1):19–25, 1988.

[21] Paul F Fransz, Susan Armstrong, J.Hans de Jong, Laurence D Parnell, Cees van Drunen, Caroline Dean, Pim Zabel, Ton Bisseling, and Gareth H Jones. Integrated Cytogenetic Map of Chromosome Arm 4S of <em>A. thaliana</em>: Structural Organization of Heterochromatic Knob and Centromere Region. *Cell*, 100(3):367–376, feb 2000.

[22] Johanna Gassler, Hugo B Brandão, Maxim Imakaev, Ilya M Flyamer, Sabrina Ladstätter, Wendy A Bickmore, Jan-Michael Peters, Leonid A Mirny, and Kikuë Tachibana. A mechanism of cohesin-dependent loop extrusion organizes zygotic genome architecture. *The EMBO journal*, 36(24):3600–3618, dec 2017.

Figure S1 *(following page)*: Differences between CAST and B6 alleles of *Prdm9*. **A:** PRDM9, DMC1 and crossover enrichment at Class 1 and Class 2 PRDM9 sites for B6 and CAST alleles. Heatmap shows log fold enrichment over 500 kb surrounding region. **B:** Normalized chromatin contact matrices, averaged across all B6 Class 1 (top) and Class 2 (bottom) binding positions. **C:** Hi-C cis/total ratio (top), compartment score (middle) and log insulation score (bottom), averaged across all PRDM9 Class 1 and 2 binding positions for B6 and CAST alleles. **D:** Cohesin subunit (top), CTCF (middle), and RNA polymerase II (RNAPII – bottom) ChIP-seq tracks, averaged across all PRDM9 Class 1 and 2 binding positions for B6 and CAST alleles. **E:** Overlap between PRDM9 Class 1 and Class 2 binding sites with meiotic cohesin subunit RAD21L sites, for CAST and B6 allele – note greater cohesin overlap with B6 allele Class 2 binding. **F:** Overlap of chromHMM epigenetic state annotations with PRDM9 Class 1 and 2 binding sites for B6 and CAST indicates similar promoter/enhancer enrichment between Class 2 sites (stronger in B6), while Class 1 for the B6 allele display strong H3K4me3 enrichment expected from PRDM9's known methyltransferase activity. Heatmap shows log fold enrichment over genome-wide mean. **G:** Number of sites overlapping each chromHMM histone annotation from mouse testis, calculated for PRDM9 B6 Class 1/2 and CAST Class 1/2 sites, as well as DSB sites and for purpose of comparison the total number of base pairs of each chromHMM annotation in the genome. **H:** Number of PRDM9 B6 Class 1 sites overlapping each chromHMM histone annotation using ES instead of testis chromHMM dataset. Lack of H3K4me3 enrichment here indicates earlier enrichment with testis-dataset is meiosis-specific. **I:** Number of H3K4me3-overlapping PRDM9 B6 and CAST sites, split by H3K4me3 regions that are testis-specific, ES-specific, and shared. **J:** Proportion of A-active vs B-inactive compartment PRDM9 B6 Class 1/2 and CAST Class 1/2 sites, as well as DSB sites. Compartments defined using ES Hi-C dataset, total genomic fraction of A-B bins shown for reference. **K:** PRDM9, DMC1 and crossover enrichment at top (most DSB favoured) and bottom PRDM9 B6 Class 1 binding loci. Heatmap shows log fold enrichment over 500 kb surrounding region. **L:** Hi-C cis/total ratio (top) and compartment score (bottom), averaged across top (most DSB favoured) and bottom PRDM9 B6 Class 1 binding loci. Trends similar to CAST allele (Fig. 3 main text). **M:** Overlap of chromHMM epigenetic state annotations with top (most DSB favoured) and bottom PRDM9 B6 Class 1 binding loci. Note greater enrichment of H3K4me3 at DSB-favoured hotspots. Heatmap shows log fold enrichment over genome-wide mean.

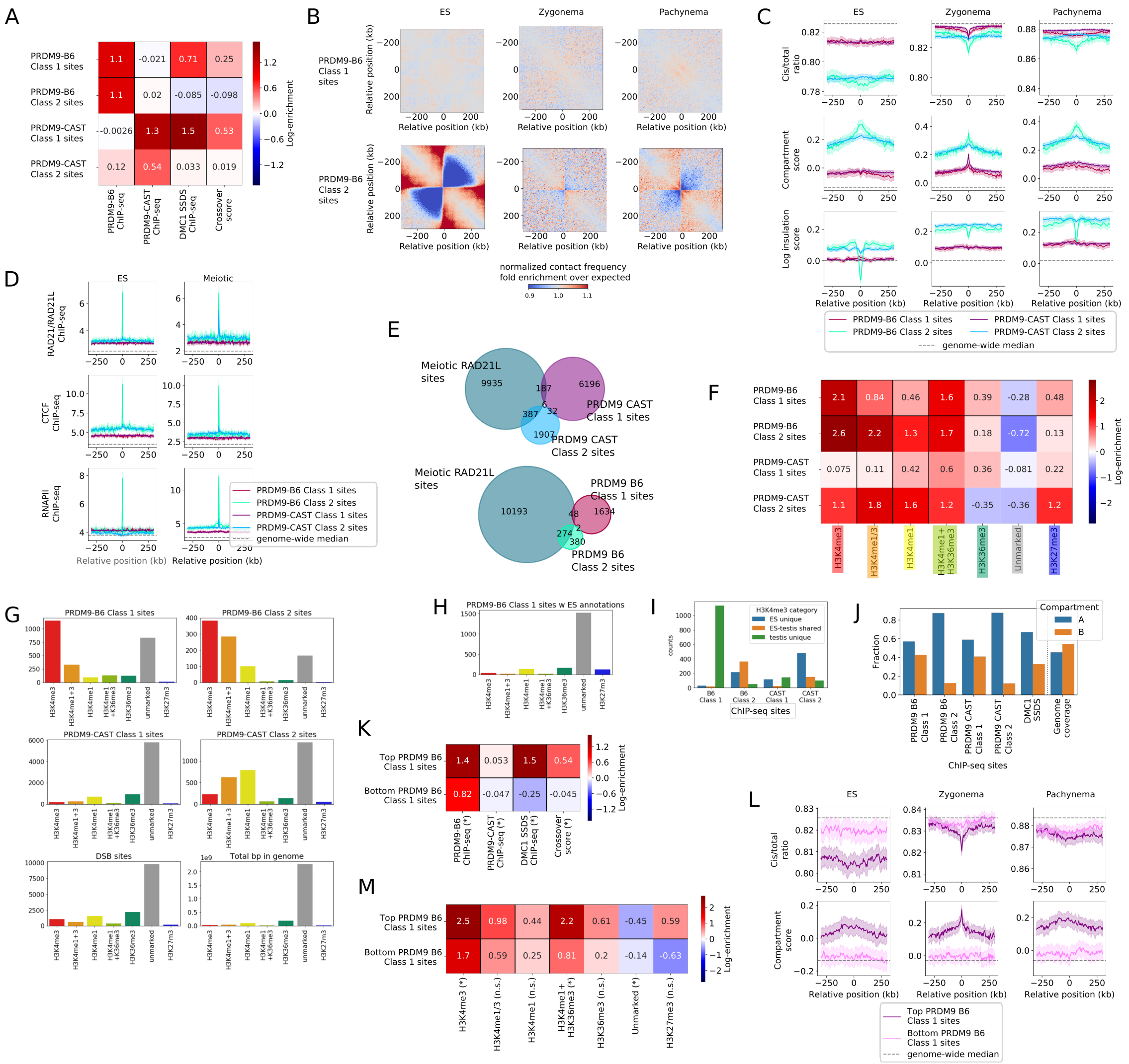
A
PRDM9-B6 Class 1 sites
PRDM9-B6 Class 2 sites
PRDM9-CAST Class 1 sites
PRDM9-CAST Class 2 sites
PRDM9-B6 ChIP-seq
PRDM9-CAST ChIP-seq
DMC1 SSDS ChIP-seq
Crossover score
Log-enrichment
B
ES
Zygonema
Pachynema
Relative position (kb)
normalized contact frequency fold enrichment over expected
C
Cis/total ratio
Compartment score
Log insulation score
PRDM9-B6 Class 1 sites
PRDM9-B6 Class 2 sites
PRDM9-CAST Class 1 sites
PRDM9-CAST Class 2 sites
genome-wide median
D
ES
Meiotic
RAD21/RAD21L ChIP-seq
CTCF ChIP-seq
RNAPII ChIP-seq
E
Meiotic RAD21L sites
PRDM9 CAST Class 1 sites
PRDM9 CAST Class 2 sites
PRDM9 B6 Class 1 sites
PRDM9 B6 Class 2 sites
F
H3K4me3
H3K4me1/3
H3K4me1
H3K4me1+ H3K36me3
H3K36me3
Unmarked
H3K27me3
G
PRDM9-B6 Class 1 sites
PRDM9-B6 Class 2 sites
PRDM9-CAST Class 1 sites
PRDM9-CAST Class 2 sites
DSB sites
Total bp in genome
H
PRDM9-B6 Class 1 sites w ES annotations
I
H3K4me3 category
ES unique
ES-testis shared
testis unique
counts
ChIP-seq sites
J
Compartment
A
B
Fraction
PRDM9 B6 Class 1
PRDM9 B6 Class 2
PRDM9 CAST Class 1
PRDM9 CAST Class 2
DMC1 SSDS
Genome coverage
ChIP-seq sites
K
Top PRDM9 B6 Class 1 sites
Bottom PRDM9 B6 Class 1 sites
PRDM9-B6 ChIP-seq (*)
PRDM9-CAST ChIP-seq (*)
DMC1 SSDS ChIP-seq (*)
Crossover score (*)
L
ES
Zygonema
Pachynema
Cis/total ratio
Compartment score
Relative position (kb)
Top PRDM9 B6 Class 1 sites
Bottom PRDM9 B6 Class 1 sites
genome-wide median
M
Top PRDM9 B6 Class 1 sites
Bottom PRDM9 B6 Class 1 sites
H3K4me3 (*)
H3K4me1/3 (n.s.)
H3K4me1 (n.s.)
H3K4me1+ H3K36me3 (*)
H3K36me3 (n.s.)
Unmarked (*)
H3K27me3 (n.s.)
Log-enrichment

Figure S2 *(following page)*: ChIP-seq occupancy patterns for cohesin subunits in relation chromosomal organization. **A:** (Left) Overlap in binding sites for two meiotic cohesin subunits REC8 and RAD21L, and interphase RAD21 cohesin subunit from ES cells. (Right) Comparison of RNAPII, CTCF and cohesin occupancy in meiosis and ES. **B:** Hi-C cis/total ratio (top), compartment score (middle) and log insulation score (bottom), averaged across RAD21, REC8 and RAD21L sites, calculated for ES, zygonema, and two pachynema datasets. Shading represents 95% confidence intervals. Meiotic REC8 and RAD21L sites are associated with clear dips in the log-insulation score across meiotic and ES datasets – dips are largely confined to the ES dataset for RAD21 sites. All cohesin sites are associated with positive A-B compartment scores, especially RAD21L. Meiotic RAD21L sites are generally associated with minima in cis/total, whereas the opposite effect is observed at RAD21 sites. However, in the Wang et al. pachynema dataset, a small but distinct positive cis/total ratio spike (black arrow) can be observed at RAD21L and REC8 sites. **C:** Overlap of chromHMM histone annotations with meiotic REC8, RAD21L and ES RAD21 sites. RAD21L exhibits strongest enrichment towards promoter-like histone marks (H3K4me3/H3K27me3, i.e. both active and inactive), and away from unmarked chromatin. Heatmap shows log fold enrichment over genome-wide mean, Insets plot overlap fraction averaged around cohesin sites, shading represents 95% confidence intervals. **D:** Cis/total ratio derived from pachynema Hi-C[1], averaged across cohesin binding sites split by chromHMM epigenetic category. Note that meiotic cohesin sites not intersecting active promoters or gene bodies (H3K4me3 / H3K36me3) display patterns of cis/total ratio maxima. **E:** Cis/total ratio averaged across RAD21, RAD21L and REC8 sites, derived from additional Hi-C datasets: lepto/zygotene, pachytene/diplotene spermatocytes[3] and round spermatids[3, 7]. Note elevated cis/total ratio at cohesin sites in these later meiotic / post-meiotic datasets datasets. **F:** (Top left) Schematic of Hi-C window averaging approach, with orange on-diagonal windows at ChIP-seq sites, and purple windows at off-diagonal site-pairs. Normalized chromatin contact matrices, averaged across meiotic RAD21L (bottom left), REC8 (top right) and ES RAD21 (bottom right) sites (orange) / bin-pairs (purple), for ES, zygonema and pachynema Hi-C datasets introduced earlier from Bonev et al. / Patel et al.[4, 1], and an additional pachynema dataset from Wang et al.[2]. During meiosis and in ES cells, RAD21L ChIP-seq sites exhibit boundary-like contact patterns, and site pairs exhibit enriched Hi-C contacts – patterns become more evident as meiosis progresses from zygonema to pachynema. RAD21 sites exhibit similar contact patterns strongly in the ES interphase Hi-C dataset, but weakly during meiosis (note different colour-scale). **G:** (Left) Averaged pachynema[1] Hi-C contact maps for RAD21L sites that uniquely intersect negative and positive strand transcription start sites (TSS). Downstream and upstream contacts are enriched for + and - strand sites respectively. (Right) Averaged pachynema[1] Hi-C contact maps for RAD21L bin-pairs with convergent and divergent TSS.

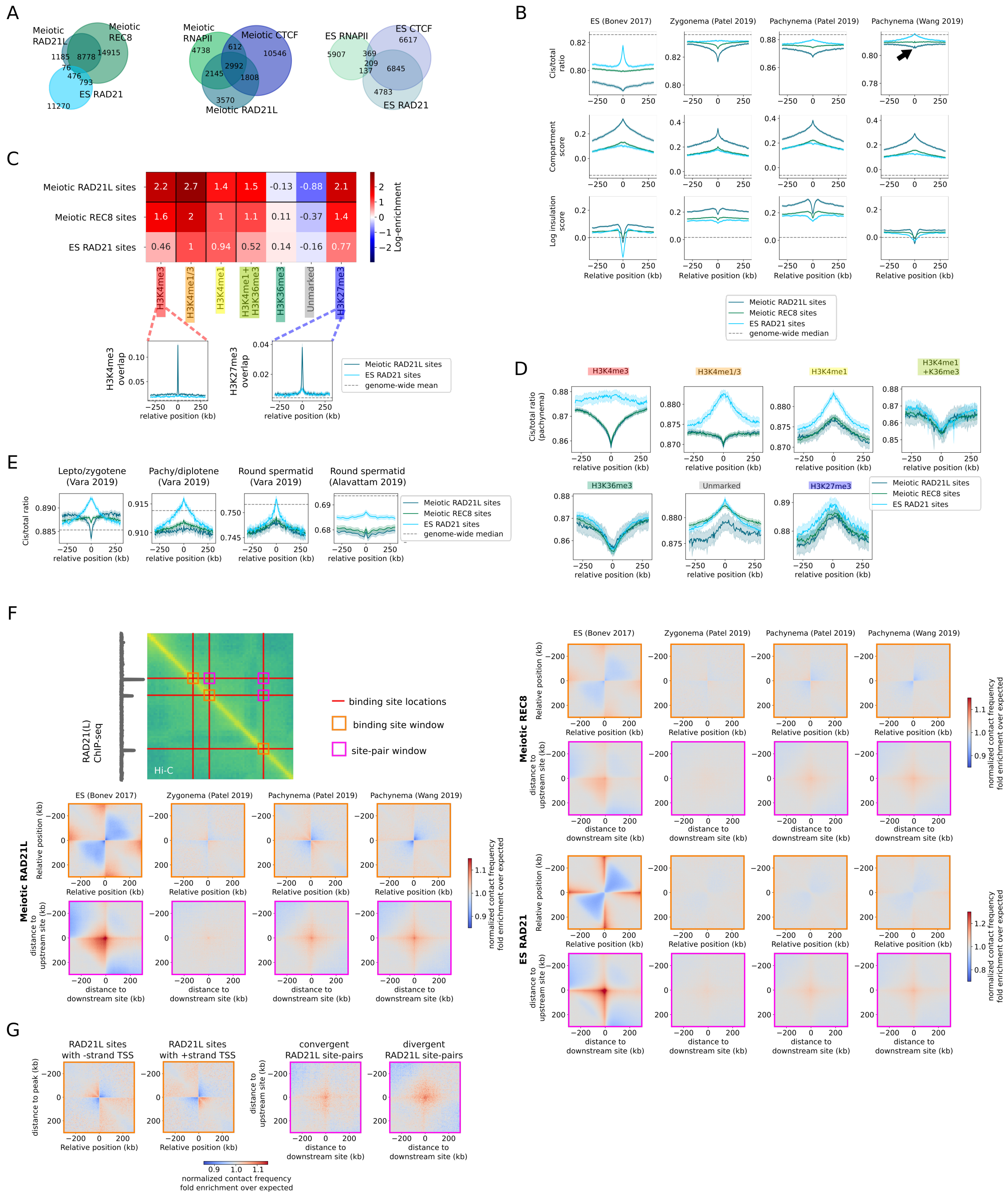
A
Meiotic RAD21L
Meiotic REC8
ES RAD21
Meiotic RNAPII
Meiotic CTCF
Meiotic RAD21L
ES RNAPII
ES CTCF
ES RAD21
B
ES (Bonev 2017)
Zygonema (Patel 2019)
Pachynema (Patel 2019)
Pachynema (Wang 2019)
Cis/total ratio
Compartment score
Log insulation score
Relative position (kb)
Meiotic RAD21L sites
Meiotic REC8 sites
ES RAD21 sites
genome-wide median
C
Meiotic RAD21L sites
Meiotic REC8 sites
ES RAD21 sites
Log-enrichment
H3K4me3
H3K4me1/3
H3K4me1
H3K4me1+H3K36me3
H3K36me3
Unmarked
H3K27me3
H3K4me3 overlap
H3K27me3 overlap
relative position (kb)
genome-wide mean
D
Cis/total ratio (pachynema)
H3K4me1+K36me3
E
Lepto/zygotene (Vara 2019)
Pachy/diplotene (Vara 2019)
Round spermatid (Vara 2019)
Round spermatid (Alavattam 2019)
Cis/total ratio
F
RAD21(L) ChIP-seq
Hi-C
binding site locations
binding site window
site-pair window
Meiotic RAD21L
Meiotic REC8
ES RAD21
distance to upstream site (kb)
distance to downstream site (kb)
normalized contact frequency fold enrichment over expected
G
RAD21L sites with -strand TSS
RAD21L sites with +strand TSS
convergent RAD21L site-pairs
divergent RAD21L site-pairs
distance to peak (kb)

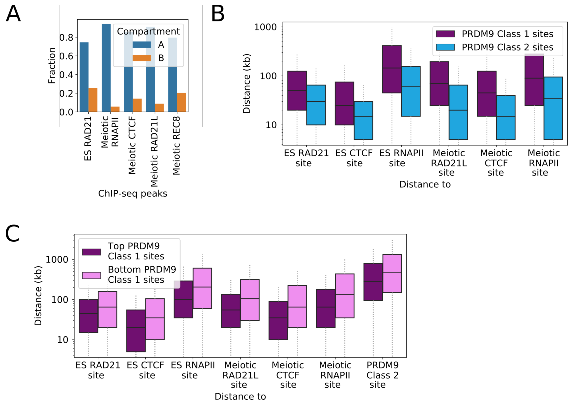


Figure S3: Cohesin and related markers are enriched in A-compartment chromatin **A:** A-B fractions of cohesin subunits and meiotic CTCF and RNAPII. **B:** As a result of clustering in A-compartment, PRDM9 Class 2 binding sites appear closer on average to cohesin and related markers compared to Class 1 sites. **C:** As a result of clustering in A-compartment, DSB favoured hotspots appear closer on average to cohesin and related markers compared to DSB-disfavoured hotspots.

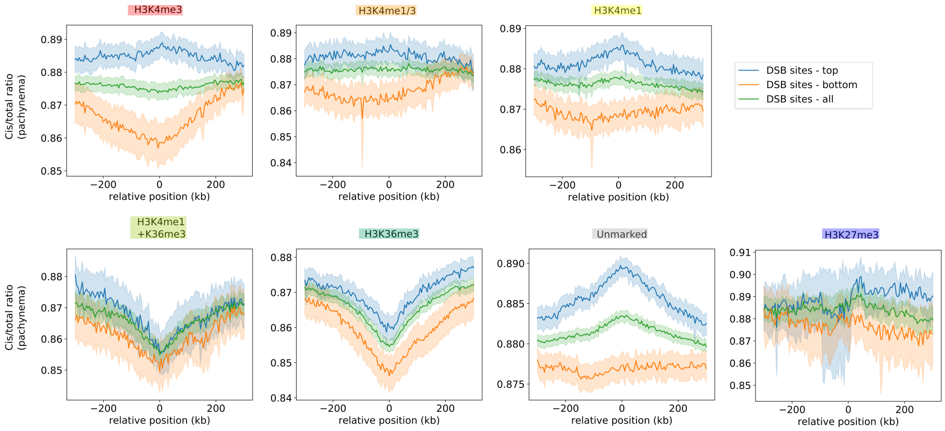


Figure S4: Pachynema cis/total ratio traces averaged across DSB sites. DSB sites were first split by chromHMM annotation into seven categories, then for each category the average of *i:* the top most crossover-likely quartile (blue), *ii:* the bottom least crossover-likely quartile (orange) and *iii:* all sites (green) were plotted.

---

Figure S5 *(following page)*: Application of principal component transformation to address collinearity between chromatin organization variables **A:** Collinearity between chromatin organization variables is demonstrated by plotting pairwise Spearman correlation coefficients between all variables. **B:** Percent explained variance for each principal component.

A

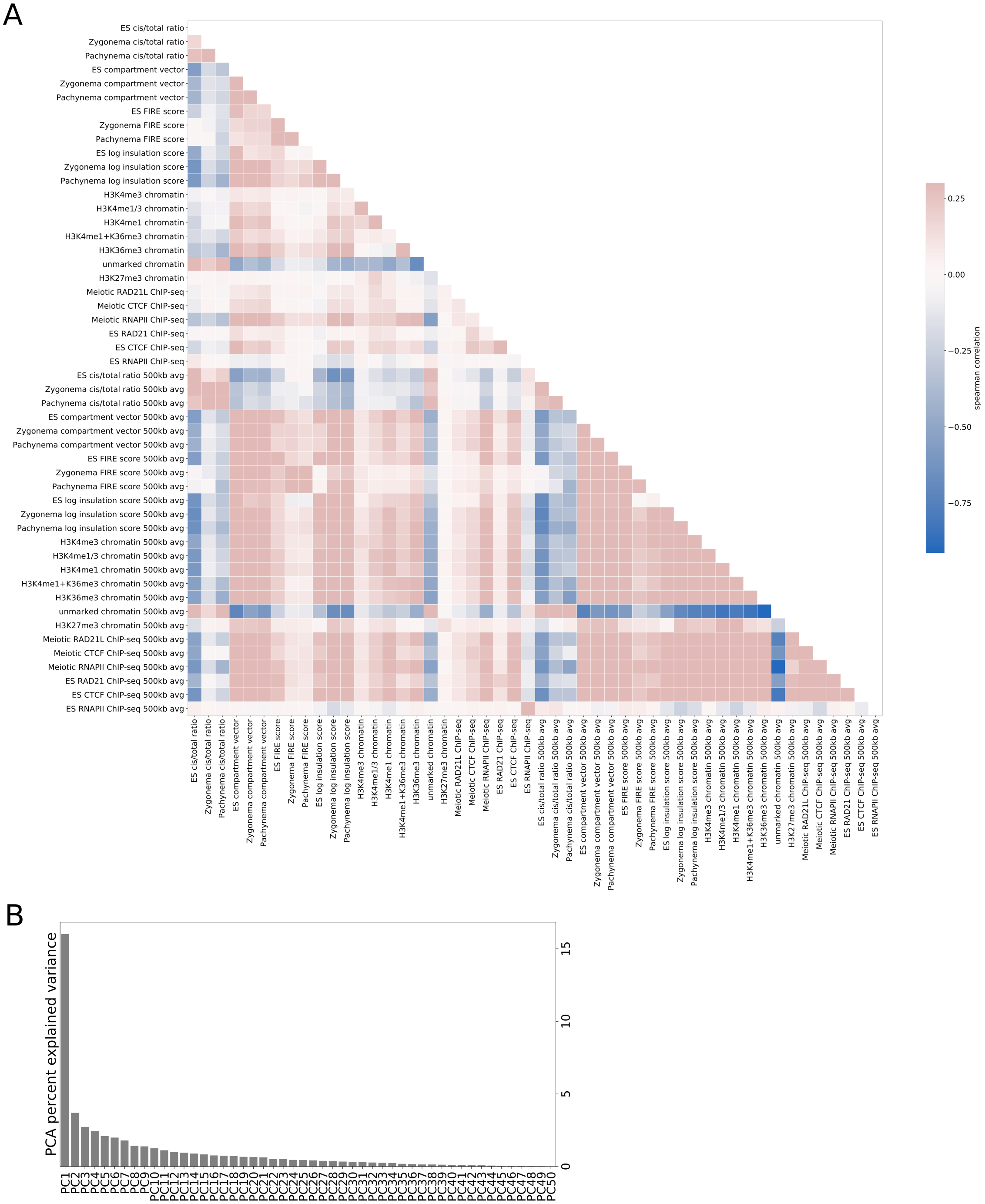
ES cis/total ratio
Zygonema cis/total ratio
Pachynema cis/total ratio
ES compartment vector
Zygonema compartment vector
Pachynema compartment vector
ES FIRE score
Zygonema FIRE score
Pachynema FIRE score
ES log insulation score
Zygonema log insulation score
Pachynema log insulation score
H3K4me3 chromatin
H3K4me1/3 chromatin
H3K4me1 chromatin
H3K4me1+K36me3 chromatin
H3K36me3 chromatin
unmarked chromatin
H3K27me3 chromatin
Meiotic RAD21L ChIP-seq
Meiotic CTCF ChIP-seq
Meiotic RNAPII ChIP-seq
ES RAD21 ChIP-seq
ES CTCF ChIP-seq
ES RNAPII ChIP-seq
ES cis/total ratio 500kb avg
Zygonema cis/total ratio 500kb avg
Pachynema cis/total ratio 500kb avg
ES compartment vector 500kb avg
Zygonema compartment vector 500kb avg
Pachynema compartment vector 500kb avg
ES FIRE score 500kb avg
Zygonema FIRE score 500kb avg
Pachynema FIRE score 500kb avg
ES log insulation score 500kb avg
Zygonema log insulation score 500kb avg
Pachynema log insulation score 500kb avg
H3K4me3 chromatin 500kb avg
H3K4me1/3 chromatin 500kb avg
H3K4me1 chromatin 500kb avg
H3K4me1+K36me3 chromatin 500kb avg
H3K36me3 chromatin 500kb avg
unmarked chromatin 500kb avg
H3K27me3 chromatin 500kb avg
Meiotic RAD21L ChIP-seq 500kb avg
Meiotic CTCF ChIP-seq 500kb avg
Meiotic RNAPII ChIP-seq 500kb avg
ES RAD21 ChIP-seq 500kb avg
ES CTCF ChIP-seq 500kb avg
ES RNAPII ChIP-seq 500kb avg
0.25
0.00
−0.25
−0.50
−0.75
spearman correlation
B
PCA percent explained variance
15
10
5
0

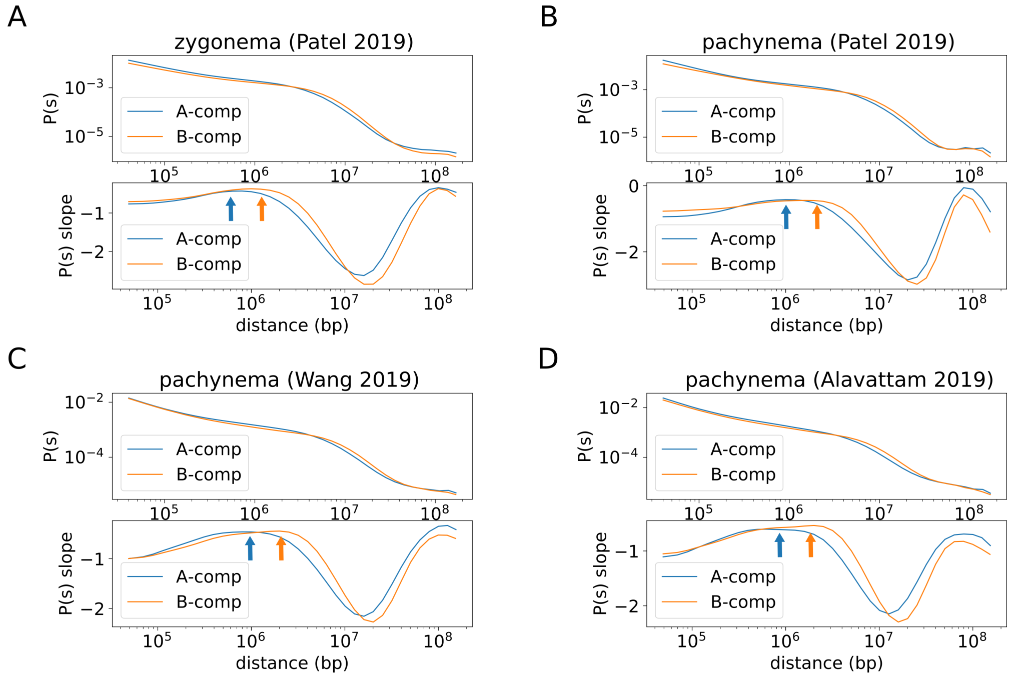


Figure S6: A-compartment loops have fewer base pairs, meaning gene-rich chromatin is closer to the chromosomal axis on average in terms of genomic distance. **A:** $P(s)$ curve derivative analysis applied to zygonema dataset from Patel *et al.* [1]. Orange and blue arrows indicate estimate of loop length for A and B compartments respectively, determined as the maxima of the derivatives of the $P(s)$ curves as in Gassler et al.[22]. **B:** $P(s)$ curve derivative analysis applied to pachynema dataset from Patel *et al.*[1] **C:** $P(s)$ curve derivative analysis applied to pachynema dataset from Wang *et al.*[2] **D:** $P(s)$ curve derivative analysis applied to pachynema dataset from Alavattam *et al.*[7]
